# Neutrophil-mediated dynamic capillary stalls in ischemic penumbra: persistent traffic jams after reperfusion contribute to injury

**DOI:** 10.1101/776427

**Authors:** Şefik Evren Erdener, Jianbo Tang, Kıvılcım Kılıç, Dmitry Postnov, John Thomas Giblin, Sreekanth Kura, I-chun Anderson Chen, Tuğberk Vayisoğlu, Sava Sakadžić, Chris B. Schaffer, David A. Boas

## Abstract

Ever since the introduction of thrombolysis and the subsequent expansion of endovascular treatments for acute ischemic stroke, it remains to be identified why the actual outcomes are less favorable despite recanalization. Here, by high spatio-temporal resolution imaging of capillary circulation in mice, we introduce the pathological phenomenon of dynamic flow stalls in cerebral capillaries, occurring persistently in the salvageable penumbra after recanalization. These stalls, which are distinct from permanent cellular plugs that can lead to no-flow, were temporarily and repetitively occurring in the capillary network, impairing the overall circulation like small focal traffic jams. In vivo microscopy in the ischemic penumbra revealed leukocytes traveling through capillary lumen or getting stuck, while red blood cell flow was being disturbed in the neighboring segments, within 3 hours after stroke onset. Stall dynamics could be modulated, by injection of an anti-Ly6G antibody specifically targeting neutrophils. By decreasing the number and duration of stalls, we were able to improve the blood flow in the penumbra within 2-24 hours after reperfusion, increase capillary oxygenation, decrease cellular damage and improve functional outcome. Thereby the dynamic microcirculatory stall phenomenon contributes to the ongoing penumbral injury and is a potential hyperacute stage mechanism adding on previous observations of detrimental effects of activated neutrophils in ischemic stroke.

**Significance:** This work provides in vivo evidence that, even in perfused capillaries, abnormal capillary flow patterns in the form of dynamic stalls can contribute to ongoing tissue injury in the salvageable penumbra in very early hours of cerebral ischemia. These events resembling micro traffic jams in a complex road network, are mediated by passage of neutrophils through the microcirculation and persist despite recanalization of the occluded artery.

## Introduction

Over the last five years, there has been a substantial expansion of endovascular recanalization practices for acute ischemic stroke with pivotal clinical trials showing evidence for functional benefit, shifting the standard-of-care for large vessel occlusions to vascular interventional methods rather than purely pharmacological(1–3). Better selection of patients with a large volume of salvageable cortical tissue with multimodal imaging technologies not only improved the success rate but has also extended the therapeutic window up to 24 hours in eligible patients(1, 4, 5). Still, however, even when eligible patients are selectively enrolled and achieve full recanalization with early intervention and when hemorrhagic complications are excluded, nearly half of these patients do not experience clinical improvement and they do not show evidence of sufficient reperfusion in brain parenchyma(6–8). This lower-than-expected efficacy has been partly attributed to subsequent arterial reocclusion and also to the “no-reflow” or “incomplete microcirculatory reperfusion” phenomenon, when microcirculatory flow is not restored in small arterioles and capillaries, despite full recanalization of the large artery(6, 9, 10). Indeed, adequacy of capillary perfusion seems to be a better predictor of clinical outcome after endovascular recanalization(6, 9, 10). Previously, sustained pericyte contraction in response to ischemia and reperfusion, possibly aggravated by reactive oxygen species, has been shown to cause red blood cell (RBC) entrapment in the ischemic tissue(11). Polymorphonuclear leukocytes have been recognized as another factor blocking the capillary lumen during the course of their activation and extravasation into the parenchyma(12–15), where they release proteases to contribute to the tissue injury. However, while these static observations are very informative, we now know that capillary flow patterns are as important as their structure for maintaining oxygen delivery into the tissue(16). This recognition motivated us to investigate the temporal dynamics of the ischemia-related persistent capillary perfusion loss, i.e. whether the capillary occlusions are permanent or they can be transient and repetitive in a particular capillary segment. Specifically, we aimed to extend our recent observations of spontaneous temporary RBC flow interruptions in cerebral capillaries, i.e. capillary stalls, in healthy animals(17) to the ischemic tissue early after recanalization and tested if neutrophils, a large and less deformable cell moving rather slowly in microvasculature(18), could be an aggravating factor the stall events across the capillary bed. By analogy, dynamic stalls occurring above critical level in the ischemic penumbra would indicate micro-level traffic jam-like formations due to extremely heterogeneous flow, as road traffic models demonstrate even tiny irregularities in vehicle flow can cause expanding flow redistributions, rapidly growing instability and subsequent accumulation of flow blocks(19, 20). We also wanted to evaluate the potential contribution of those microcirculatory abnormalities to the ongoing damage in the dysfunctional and critically hypoperfused but salvageable tissue, the ischemic penumbra(10), rather than being a bystander phenomenon. To address these questions, we took advantage of *in vivo* microscopic imaging methods, such as optical coherence tomography (OCT) and two-photon microscopy (TPM), using a mouse model of transient distal middle cerebral artery occlusion (dMCAO). The dynamic pathological perspective that we bring with those experiments, can help us understand the similar blood cell passage-related problems in other acute and chronic neurologic pathologies, potentially allowing novel therapeutic approaches.

## Results

### Mapping the Salvageable Ischemic Penumbra with OCT

We utilized a mouse model of transient dMCAO by applying mechanical compression with a device made by fusing two glass micropipettes with blunted tips (Fig1/a-c), with slight modifications of a previously reported model(23, 37, 38) that ensured stable occlusions for 60 minutes. This technique also avoided nonspecific intracranial pressure effects as shown by sham experiments (see below). Complete recanalization of dMCA was achieved simply by release of the compression after 1 hour, without any requirement for a thrombolytic agent(23, 38). Cerebral blood flow was continuously monitored during experiments by real-time laser speckle contrast imaging to ensure successful occlusion and recanalization (Supplementary Movies 1 and 2, Fig2/a-c). In our experience, the success rate for achieving a sufficient level of occlusion was around 95%. The occlusion was considered successful if at least one pial MCA segment within the cranial window had absence of blood flow, in addition to opening of collaterals from the anterior cerebral artery (ACA)-supplied zone (Supplementary movie 1). Another criterion for successful occlusion was appearance of abnormal optical scattering measured by OCT within the cranial window (see below). Recanalization was successful in all experiments (as demonstrated by recovery of blood flow in pial MCA branches) with no visible residual damage to the dMCA (Fig1/e). Since all animals had different pial vascular anatomy and level of collateral supply, and we wanted to study the abnormal capillary dynamics in the penumbra tissue immediately next to the evolving core in every single experiment, we had to control for variations that could result in different spatial distributions of varying grades of oligemia. Real-time monitoring of blood flow changes by laser speckle contrast imaging allowed us to assess the baseline flow distribution and postischemic collateral flow and to guide the subsequent OCT imaging of microcirculation to the penumbral zones supported by the collaterals. Longitudinal OCT imaging within this area at baseline and different time point during vascular occlusion/recanalization (Fig 1/d) was used to assess cerebral blood flow quantitively, to visualize capillary flow and optical attenuation changes. As Precise differentiation between core and penumbra was not readily apparent with laser speckle contrast imaging alone (Fig2/a-c), OCT imaging data properly mapped the infarct area inside the MCA-supplied ischemic cortex, helping us direct our further analyses to the core-penumbra transition zones on an individualized basis. Our dMCAO model also allowed us to conduct the experiment without removing or repositioning the animal under the imaging equipment; an advantage over proximal MCAO models.

**Figure 1.**
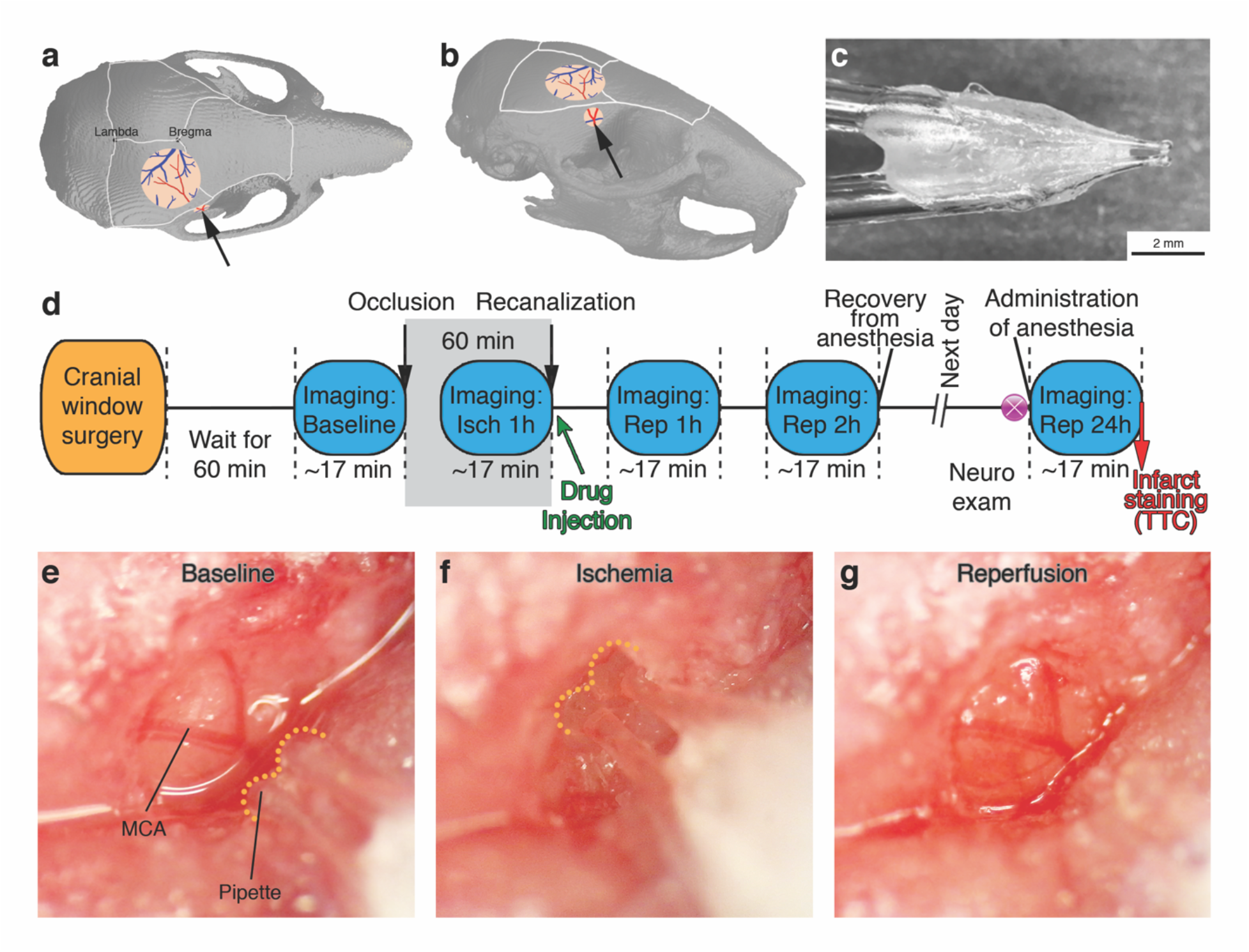
**(a,b)** Diagrams showing the top and oblique views of the mouse skull, with the locations of two cranial windows indicated. Black arrow shows the direction of compression over the distal MCA at the bifurcation point. **(c)** Close-up view of the occlusion device, made by fusion of two blunted micropipettes with an approximate 10-degree angle at the tip. **(d)** Experimental timeline showing the surgery, MCA occlusion and recanalization, drug injection imaging time points and study end points. **(e-g)** Color photographs acquired under the surgical microscope, showing the experimental procedure of dMCAO and recanalization. The tip of the micropipette device is outlined with yellow dotted line. After recanalization there was no visible injury to the MCA, apart from minimal bruising in the dura. A reactive dilatation in dMCA can be noted.

**Figure 2.**
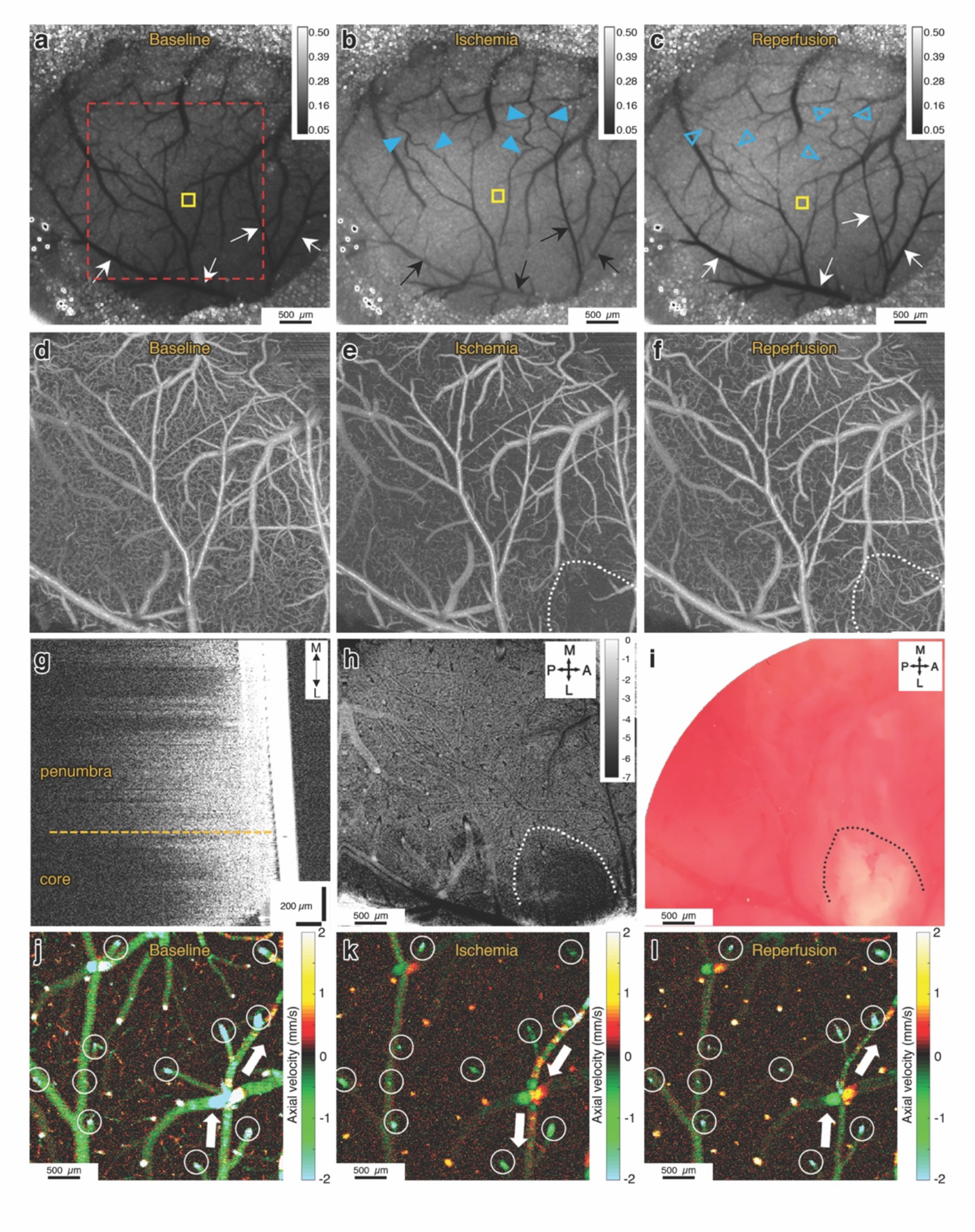
**(a-c)** Laser speckle contrast images showing the baseline, ischemia (end of 1^st^ hour) and recanalization (after 1 hour) conditions. Black and white arrows show the pial branches of the MCA. Darker colors indicate higher flow. After occlusion, we see decreased or absent flow in the pial MCA branches, and appearance of flow in collateral vessels (blue arrowheads) from the ACA-supplied region. After recanalization of dMCA, we see collateral flow disappears (blue empty arrowheads). Scale bars: 500 µm. **(d-f)** OCT angiograms (average of 10 images) of the area shown in the red square in panel (a). Only RBC-perfused capillaries are visible in those images. During ischemia, there is a decreased number of perfused capillaries in the imaged area, but more severely in the outlined ischemic core (inside the white dotted line). One hour after recanalization, peripheral penumbra zone has sustained low capillary flow. Scale bars: 500 µm. **(g)** Sagittal cross-section showing the axial decay of OCT amplitude signal. Solid white structure on the right is the cover glass. There is reduced axial penetration of the OCT signal in the core area. **(h)** Signal attenuation slope map of the same region imaged simultaneously with (f). Darker pixels indicate higher (more negative) axial signal attenuation slope. Unit of calibration bar is 1/mm^2^. **(i)** The histological infarct shown by lack-of TTC staining of the brain approximately 10 min after the imaging in (i) shows overlap with the zone of reduced signal penetration. **(j-l)** prD-OCT images of axial velocity in the ischemic penumbra, during baseline, ischemia and recanalization. Imaging planes are approximately 50 µm below the cortical surface, to get optimal cross-sections of penetrating arterioles. Arterioles and venules could be differentiated by their flow direction, as indicated by different colors. Negative values indicate arterioles and positive values indicate venules. Blood flow values were measured longitudinally in each penetrating arteriole (white circles) for quantification across time points. White arrows show the direction of flow in a distal pial branch of MCA, that was reversed, receiving collateral flow during occlusion and resolved after recanalization. Scale bars: 100 µm.

Approximately 60 min after the initiation of dMCAO, OCT angiograms demonstrated a severely compromised microvascular bed with almost no capillary perfusion in the ischemic core and relatively better, but still lower-than-baseline perfused capillary density in the penumbra region, surrounding the ischemic core (Fig 2/d-e). We also detected an abnormal optical scattering within the ischemic core, allowing a clear distinction of the developing infarct within the hypoperfused MCA territory. To quantify the scattering changes, the axial decay profile of the OCT signal amplitude was analyzed as a function of cortical depth by following the procedure outlined in a previous study(39). The axial signal attenuation was prominently higher (i.e. had a higher axial decay slope) in the well-demarcated area with very low (or absent) capillary perfusion (the core) compared to the attenuation in the periinfarct penumbral zones (Fig 2/l-m, Supplementary Figure 2). These attenuation changes established 2 hours after recanalization overlapped with the infarcts demonstrated by TTC staining done shortly after in vivo imaging (Fig2/n). Modeling the well-documented MRI-definition of diffusion-perfusion mismatch(40), we identified our penumbral region of interest as the hypoperfused MCA region peripheral to the ischemic core border. The change in the slope of the axial OCT signal in the evolving infarct was not a direct result of cerebral blood flow and/or blood volume changes as the attenuation changes did not synchronize with loss of capillary flow (Supplementary figure 3).

Phase-Resolved Doppler OCT (prD-OCT) was used to quantify flow velocity in individual penetrating arterioles under baseline and ischemic conditions, allowing us to calculate the cerebral blood flow in a given region of interest. Because of baseline variability in cerebral blood flow, we expressed the subsequent ischemia and reperfusion values as normalized to baseline. This measure was used to compare blood flow changes in different experimental groups, as detailed below. In overall, within the ischemic core, blood flow was found to be decreased to 14±4% of baseline during MCAO, while in penumbra it decreased to 39±4% and these values were in agreement with previously reported levels in similar models of cerebral ischemia(41, 42).

One hour after MCA recanalization, we consistently detected a persisting lower-than baseline capillary perfusion in the ischemic penumbra. These changes were not confounded by systemic physiological parameters, like oxygen saturation, heart rate, blood pressure and body temperature (Table 1). In the next step, we wanted to further evaluate the capillary-level flow dynamics in the ischemic penumbra, to understand if the proposed stalls were contributing to an impairment in overall blood flow.

**Table 1.**
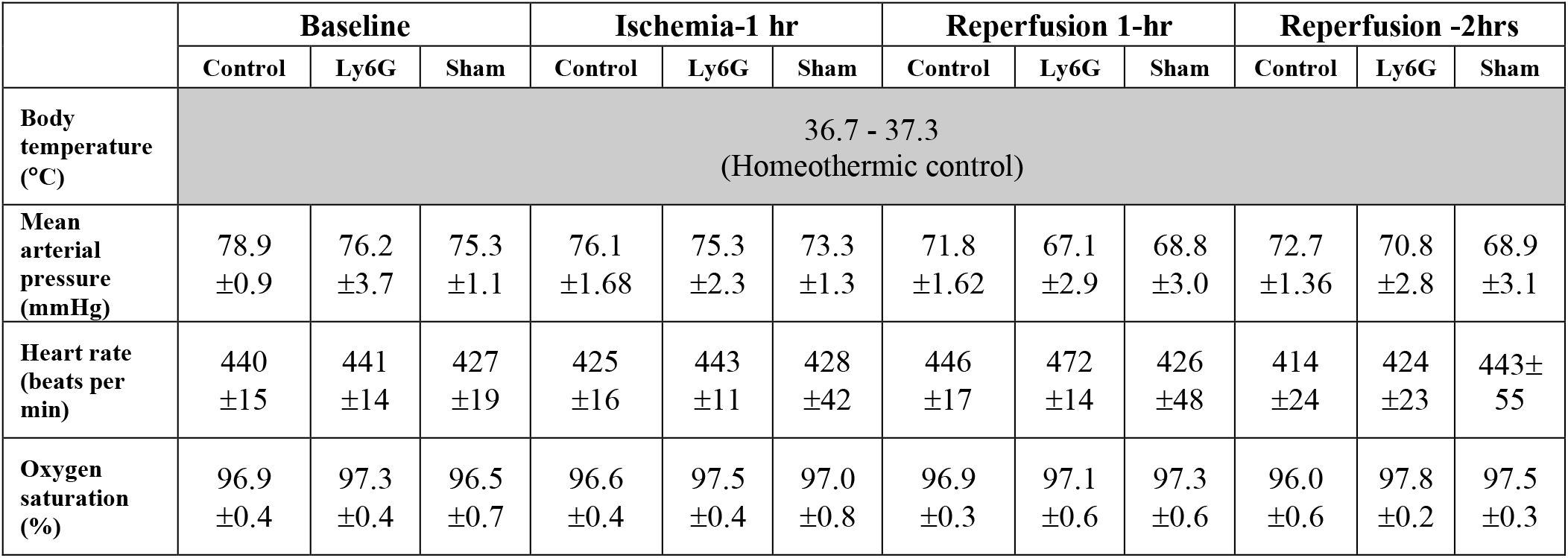
Physiological parameters.

|  | Baseline |  |  | Ischemia-1 hr |  |  | Reperfusion 1-hr |  |  | Reperfusion -2hrs |  |  |
| --- | --- | --- | --- | --- | --- | --- | --- | --- | --- | --- | --- | --- |
|  | Control | Ly6G | Sham | Control | Ly6G | Sham | Control | Ly6G | Sham | Control | Ly6G | Sham |
| Body temperature (°C) | 36.7 - 37.3<br>(Homeothermic control) |  |  |  |  |  |  |  |  |  |  |  |
| Mean arterial pressure (mmHg) | 78.9<br>$\pm 0.9$ | 76.2<br>$\pm 3.7$ | 75.3<br>$\pm 1.1$ | 76.1<br>$\pm 1.68$ | 75.3<br>$\pm 2.3$ | 73.3<br>$\pm 1.3$ | 71.8<br>$\pm 1.62$ | 67.1<br>$\pm 2.9$ | 68.8<br>$\pm 3.0$ | 72.7<br>$\pm 1.36$ | 70.8<br>$\pm 2.8$ | 68.9<br>$\pm 3.1$ |
| Heart rate (beats per min) | 440<br>$\pm 15$ | 441<br>$\pm 14$ | 427<br>$\pm 19$ | 425<br>$\pm 16$ | 443<br>$\pm 11$ | 428<br>$\pm 42$ | 446<br>$\pm 17$ | 472<br>$\pm 14$ | 426<br>$\pm 48$ | 414<br>$\pm 24$ | 424<br>$\pm 23$ | 443±<br>55 |
| Oxygen saturation (%) | 96.9<br>$\pm 0.4$ | 97.3<br>$\pm 0.4$ | 96.5<br>$\pm 0.7$ | 96.6<br>$\pm 0.4$ | 97.5<br>$\pm 0.4$ | 97.0<br>$\pm 0.8$ | 96.9<br>$\pm 0.3$ | 97.1<br>$\pm 0.6$ | 97.3<br>$\pm 0.6$ | 96.0<br>$\pm 0.6$ | 97.8<br>$\pm 0.2$ | 97.5<br>$\pm 0.3$ |

### Dynamic Capillary Stalls Increase in Ischemic Penumbra and Persist After Reperfusion

In our preliminary experiments, OCT angiogram time series over ∼6.5 min consistently showed very frequent transient RBC stalls (compared to our experience in healthy animals(17)) in individual capillaries during ischemia (Fig3/a-g) and for at least 2 hours after reperfusion. Interestingly, this dynamic information was lost if capillary angiograms were averaged, as commonly done with OCT angiogram data to increase the signal-to-noise ratio (Supplementary movie 3). The minimum duration of a stall that we could detect in our experiments, limited by the OCT frame rate, was 6.5 seconds. The main advantage of OCT with this evaluation was its ability to image the flow in several hundred capillaries simultaneously and the very prominent signal change when capillary RBC flow stops. Two photon fluorescent angiography in the corresponding areas with frequent and dynamic stalls 2 hours after reperfusion showed repetitive plugging of capillaries by cells identified as leukocytes based on their size (Supplementary movies 5 and 6). We further confirmed these cells were leukocytes but not RBC clusters by staining them with Rhodamine-6G, a mitochondrial stain (Supplementary movie 7, Supplementary figure 4). These leukocytes would plug individual capillaries for seconds or minutes, then would squeeze slowly through the occluded capillary and finally allow flow in that particular capillary again. A single temporary plug could affect the RBC flow in other capillaries in the nearby network, dynamically affecting the propensity of stalls in a greater number of capillaries (Supplementary movie-6). We hypothesized that capillary flow irregularities introduced by stuck or slowly moving leukocytes would be contributing to the dynamic persistent capillary stalls in the reperfused ischemic penumbra, as if they were slowly moving trucks in a busy traffic. We were also motivated by a recent study on increased number of capillary flow stalls in a mouse model of Alzheimer’s disease(43) that could be resolved by specific targeting of a neutrophil surface protein, Ly6G, using a monoclonal antibody, resulting in improved cerebral blood flow and cognitive performance within minutes. Therefore, we aimed to improve the dynamic persistent capillary flow interruptions in the penumbra after ischemia-reperfusion by targeting neutrophils in our transient stroke model with the same type and dose of monoclonal antibody, comparing the effect against isotype-control antibodies. Although our primary aim was to identify at least one cellular role player in the increased number of stalls rather than establishing a treatment method with full potential, we also wanted to briefly test the possibility of using anti-Ly6G as an adjunctive target for endovascular recanalization of a stroke patient. Therefore, we chose to apply the antibody intravenously (2mg/kg) immediately after recanalization of the MCA (n=11). We used that specific dose of antibody because it was found effective for improving leukocyte-dependent stall in the previous study(43). For controls, the same dose and volume of isotype control antibodies were injected (n=13).

**Figure 3.**
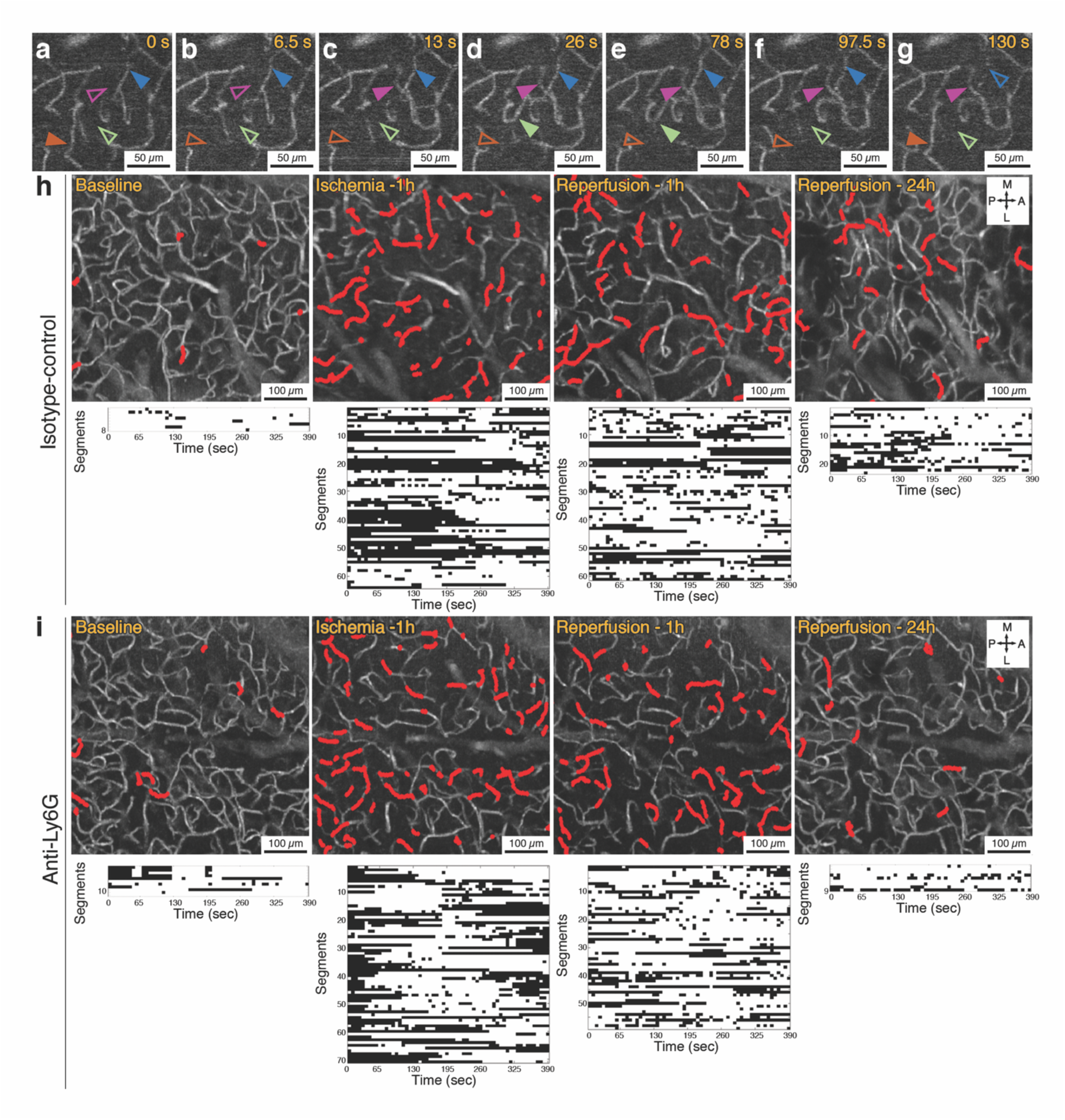
**(a-g)** Representative OCT angiogram time series showing capillary flow appearing and disappearing repeatedly over an observation period of 130 seconds in the penumbra, during one hour of MCA occlusion. Solid arrowheads indicate that the capillary segment is flowing and empty arrowheads indicate that the segment is not flowing at that time point. Scale bars: 50 µm. **(h)** Capillary angiograms over penumbra zone of an isotype-control injected animal (average of 60 images, acquired over 6.5 min) with the dynamically stalling segments indicated in red. Each stall event measured in these segments are shown in the plots (the bottom row) as black dots. The extremely high stall fraction and rate was apparent during occlusion but also persisted after reperfusion. **(i)** In an animal injected with anti-Ly6G antibody immediately after reperfusion, although stall profile during baseline and dMCAO were similar, there was profoundly better capillary flow profile at 24 hours. Scale bars in (h) and (i): 100 µm.

In both Ly6G-targeted and control groups, one hour after induction of ischemia by dMCA compression, the incidence of stalling capillaries increased almost threefold and the prevalence almost six-fold, indicating that nearly 20% of the visible capillaries (i.e. capillaries exhibiting flowing red blood cells at some point during our 6.5 min measurement) were not flowing in the ischemic penumbra at any given time, with a different subset of capillaries flowing at each time point (Fig3/h and Fig 4/d-f). This observed effect on dynamic stalls was proportionally higher than the 50% reduction in perfused capillaries at any time (Fig4/c) (this percentage of capillary perfusion reduction was also in line with previous studies(44)). Any contributing effect of a coincident peri-infarct depolarization to the dynamic capillary occlusions was excluded by continuously monitoring the blood flow with laser speckle contrast imaging; since these waves are accompanied by spreading oligemic waves(45). Moreover, the capillary stalls were happening without observable fluctuations of flow in large blood vessels, hence being an exclusive phenomenon of the microcirculation (supplementary movie 3 and 4). For the subsequent two hours after MCA recanalization, stall incidence, prevalence and cumulative duration, remained statistically higher than baseline condition in the penumbra (Fig 3/h, Fig 4/d-f). This implied that even the OCT-visible capillaries in ischemic penumbra in averaged angiograms, were not flowing optimally but were showing high temporal heterogeneity per each segment. In some segments in penumbra, 2 hours after canalization, RBC flow was stopping for more than 5 min, then restarting before another interruption settled in (Fig 3/h).

**Figure 4.**
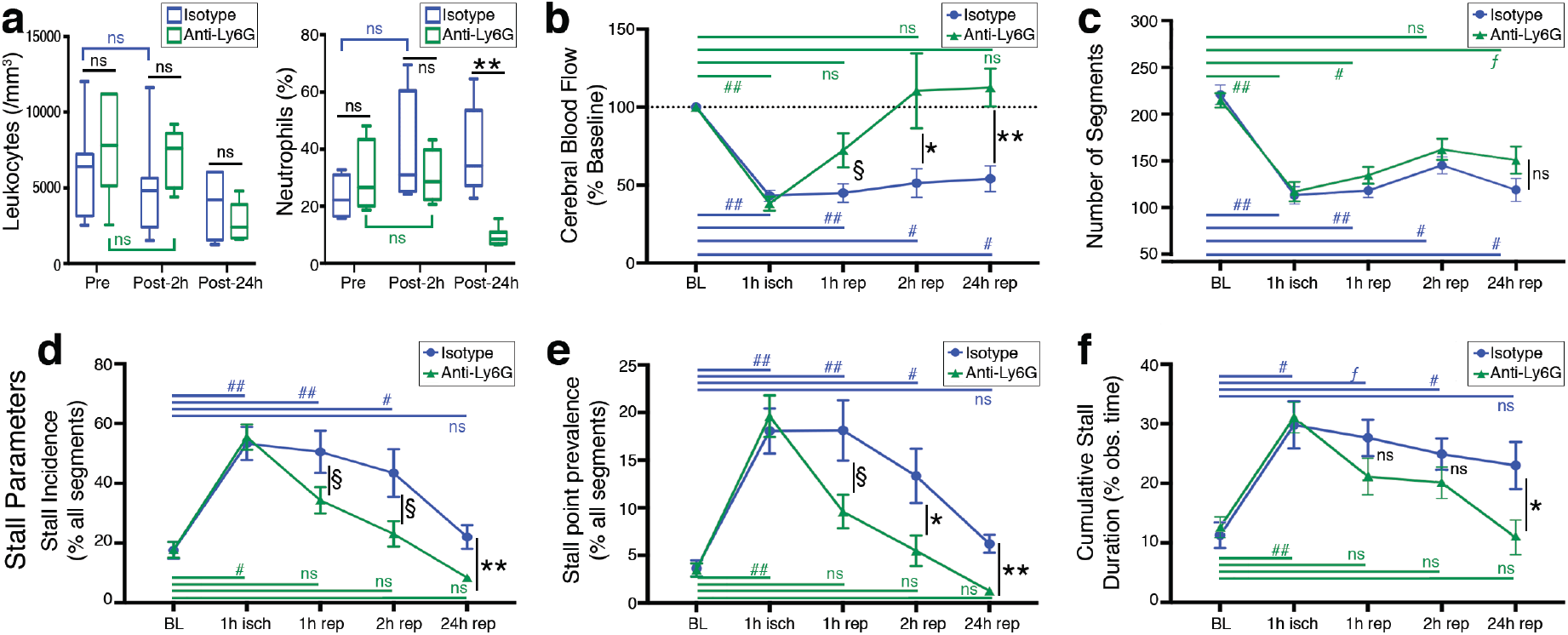
**(a)** Total leukocyte count was not significantly different in Ly6G group from control at any time point, despite being relatively lower at 24 hours. Differential neutrophil count was significantly different between two groups only at the 24-hour time point, which is consistent with a neutrophil-depletion effect of the Ly6G antibody. In control animals, there was a trend of increasing neutrophil counts after ischemia over 1 day although statistically nonsignificant. Boxes show medians and interquartile range, whiskers show 5-95^th^ percentiles. **(b)** Penumbral blood flow changes during MCAO and recanalization in control and Ly6G groups, measured by prD-OCT and expressed as percentage of baseline flow. The decrease in blood flow was almost the same in both treatment groups, but blood flow recovered to a significantly higher level at 2 and 24 hours after recanalization in the Ly6G-treated group **(c)** Total number of capillary segments in experimental groups in ischemic penumbra, measured at baseline and at different experimental time points. There were similar number of segments at baseline in both groups and ischemia significantly diminished the number of capillary segments flowing at any time point during observation, identically in both groups. The capillary count never returned to the baseline in both groups, during 1 day of observation after MCA recanalization. Although the anti-Ly6G group seemed to have a relatively higher number of capillary segments after reperfusion, this difference was not significant. **(d-f)** Longitudinal monitoring of various stall parameters at different time points during occlusion and recanalization in ischemic penumbra. Baseline and ischemia conditions were almost identical in control and Ly6G groups, but with an improvement starting (but marginally significant) as early as 1 hour, which became significant at 2 hours and 24 hours after reperfusion. The change in cumulative stall duration (g) was less obvious, but still significant at 24 hours. **Significance:** (**) (##) p<0.01, (*)(#) 0.01≤p<0.05, (ƒ)(§) 0.05≤p<0.09, (n.s.) p ≥0.09. **Sample sizes:** (a) n=6 (b-f) n=6 for 24 hours, n=8-13 for other time points. Data is shown as mean±SEM except for (a-b).

Meanwhile, we did not detect time-dependent increases in capillary stall measurements in sham animals (n=4) with nonspecific mechanical compression (that did not target the dMCA) over a 3-hour observation period (Supplementary figure 1), making a confounding effect of craniotomy or systemic phenomenon unlikely for our observations on stalls with dMCAO.

### Targeting Neutrophils by Anti-Ly6G monoclonal antibody reduces the stall events in penumbra

Ly6G is a glycosylphosphatidylinositol-anchored surface antigen specifically expressed on neutrophils, with unknown function and endogenous ligand(46, 47). This protein is particularly useful for identification of neutrophils with flow cytometry and also it helps for in vivo targeting of neutrophils to induce neutropenia in mice(47, 48). Although there was no significant difference between the two groups in baseline or ischemia (1^st^ hour) stall parameters (therefore the two groups were identical before the antibody administration), we observed a reduction in capillary stalls after recanalization in the Ly6G administered group, starting possibly at 1 hour but becoming more apparent by 2 hours (Fig 3/i, Fig4/d-f). When cranial windows were imaged at 24 hours, the difference in stalls was even more significant (Fig 3/i, Fig 4/d-f). An improvement in cerebral blood flow profile accompanied the reduction of capillary stalls, as measured by prD-OCT (Fig 4/b). It should be noted that the number of visible capillaries was not significantly different between the two treatment groups at any time point. This improvement was also independent of systemic physiological changes with no significant difference across groups at all time points (Table 1). Since anti-Ly6G treatment can lead to a depletion of neutrophils over 24 hours(47, 49), we also took blood samples to evaluate leukocyte and differential neutrophil counts before and after antibody administration. The leukocyte and differential neutrophil counts at 2 hours after reperfusion and anti-Ly6G treatment was not different from the controls (Fig 4/a), which is also in line with the previous study on Alzheimer’s models with the same monoclonal antibody(43). At the 24^th^ hour however, we saw a prominent reduction in neutrophil count compared to the control antibody treated group as expected, confirming the specific effect of the anti-Ly6G treatment on neutrophils (Fig 4/a). The modulatory effect on the microcirculation at the 1^st^ day therefore could be attributed to a reduction in neutrophil count. However, at the very acute phase as early as 2 hours, without any effect in neutrophil counts, we still noticed a significant effect on capillary flow, suggesting that reduction of neutrophil count may not be necessary for the effect to occur.

### Improving Microcirculation via Anti-Ly6G Improves Functional Outcomes and Reduces Ischemic Damage

In addition to quantifying an improvement in microcirculatory parameters after reperfusion with intravenous administration of anti-Ly6G antibody, we further tested if this improvement would be actually beneficial for the ischemic tissue, contributing to the salvaging of penumbra. In a separate group of mice, we first compared the effect of anti-Ly6G antibody (n=3) versus control (n=3) on capillary pO_2_ distribution in the acute setting of reperfusion, 2 to 4 hours after recanalization was achieved, with phosphorescence lifetime imaging (Fig 5/a-b). Excessive temporal fluctuations in capillary oxygen pressure in the presence of stalls (between 18-65 mmHg over ∼2.5 min) (Figure 5/d) was suggestive for pathological effects of stalls on oxygenation of ischemic microvasculature. And we indeed detected a significantly higher fraction of well-oxygenated capillaries with anti-Ly6G treatment in ischemic penumbra 2 hours after reperfusion (Figure 5/c).

**Figure 5.**
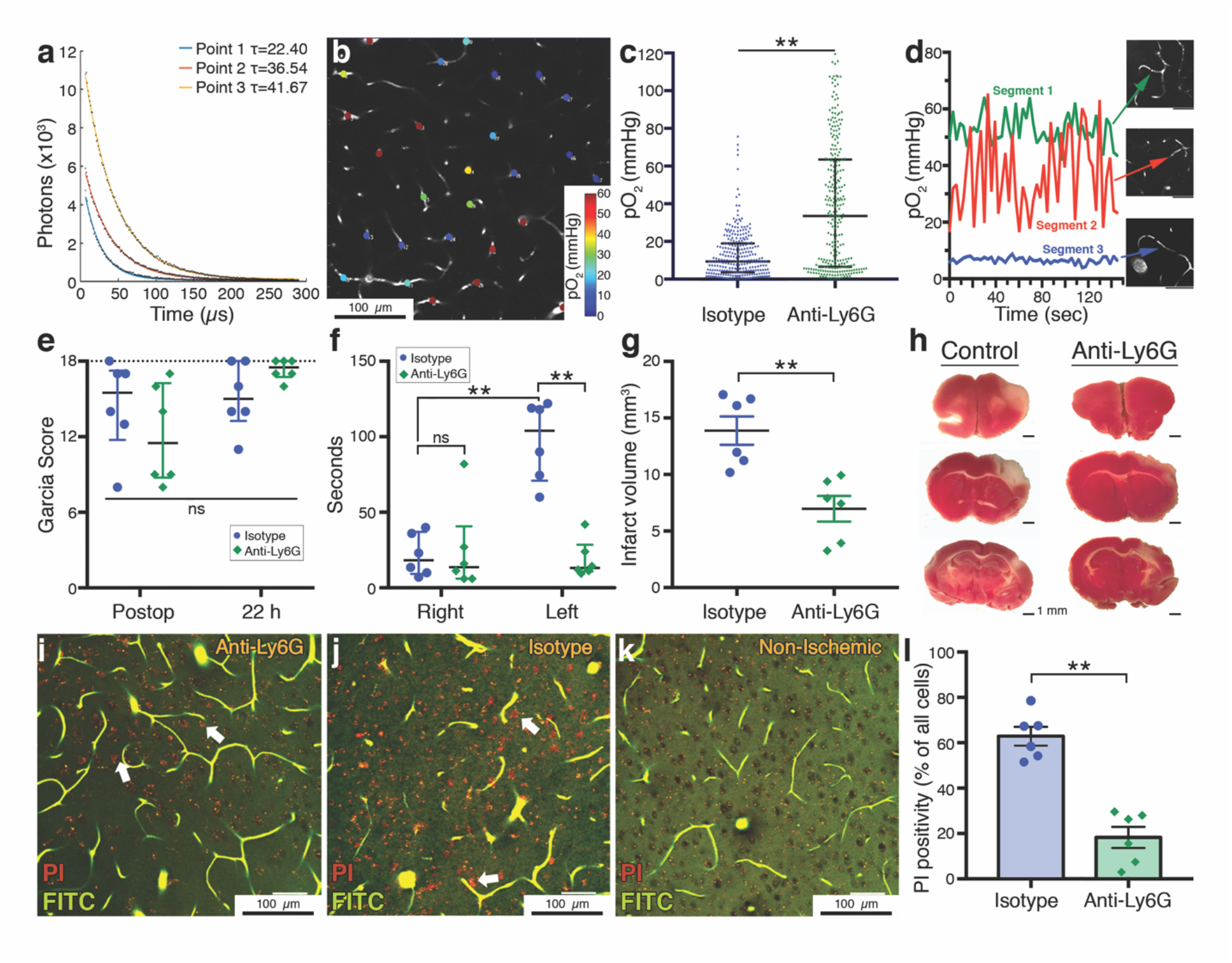
**(a)** Phosphorescence lifetime measurements at three different points with different decay time constants (τ’s) corresponding to different pO_2_ values. **(b)** Two-photon imaging plane showing pO_2_ values at measurement points across the microvasculature. Scalebar: 100 µm. **(c)** Distribution of pO_2_ values in measured capillaries in control and Ly6G-treated mice (n=3 per group) showing that Ly6G-treated mouse had a significantly higher level of capillary oxygenation. Data summarized as median-interquartile range. **(d)** Temporal pO_2_ tracings measured at 3 different capillaries, one with constant flow (segment 1), one with constant stall during the observed time (segment 3) and one with repetitive short-duration stalls (segment 2). pO_2_ partial pressure fluctuates prominently in the repeatedly stalling capillary. Scalebars: 50 µm. **(e)** Garcia neurological scores in the postoperative period, 1 hour after recovery from anesthesia was similar in both treatment groups. At 22 hours, anti-Ly6G treated animals had relatively higher scores but the difference was not significant, pointing to the relatively poor sensitivity of the Garcia score for evaluation of small infarcts generated by dMCAO. Maximum possible score of 18 is indicated with the dashed line. Data summarized as median-interquartile range **(f)** Adhesive tape removal test indicated a significant impairment in control animals compared to the right (unaffected) side, as shown here as time to removal of the tape in seconds. This asymmetry disappeared in anti-Ly6G treated animals. Data shown as median-interquartile range **(g-h)** Infarct volumes were significantly lower in Ly6G-treated animals. The cortical area with lack of pink staining demonstrates the irreversibly injured tissue. Data summarized as mean±SEM **(i-k)** Ex vivo angiograms with red propidium iodide (PI) labeling of injured cells (white arrows) in ischemic periinfarct tissue. These images are maximum intensity projections over 30 µm, at approximately 150 µm depth under cortical surface. Capillaries are visible as they are filled with a fluorescent gel containing FITC (green). Anti-Ly6G treated animals had less dense PI labeling compared to controls. No specific PI labeling was observed in nonischemic brain areas (k). Punctate red fluorescence signals originate from lipofuscin granules that are normally found in neurons. Total number of cells were determined by counting of the black circular structures visible in the relatively brighter background on green channel. Scale bars: 50 µm. **(l)** PI positivity, as expressed by the percentage of all visible cells was significantly lower in anti-Ly6G treated group. Data summarized as mean±SEM. **Significance:** (**) p<0.01, (*) 0.01 ≤p<0.05, (n.s.) p≥0.05. **Sample sizes:** n=6 mice for (e-g), n=6 regions over 3 mice per group in (j), n=80-86 vessels (per animal) for six mice in (c).

We then evaluated functional neurological scores at approximately 22 hours after recanalization (n=6 per group). Initial Garcia neurological scores of animals that were recorded 1 hour after recovery were not significantly different between the two groups and with prominent neurological deficits, suggesting that the initial ischemic insult was comparable between groups (Fig5/e). We saw an improvement indicating functional recovery with the Garcia scores at 22-hours for both groups, while the difference was not significantly different between control and anti-Ly6G-treated animals (Fig5/e), but we explained this by the relative insensitivity of this scoring system for smaller cortical lesions generated by distal MCA ischemia models(33). To account for fine sensorimotor disturbances, we performed an adhesive tape removal test, a sensitive approach suitable for small ischemic lesions generated by distal MCAO models(34). This test, indeed, revealed a significant delay in tape removal score in the affected limb of the control animals (Figure 5/f). In the anti-Ly6G-treated group, the difference between the affected and nonaffected limb was significantly decreased, showing a prominent improvement in functional state. Moreover, when brains of these animals were immediately removed approximately 24 hours after the ischemic insult and infarct volumes were determined by 2,3,4-triphenyltetrazolium chloride (TTC) staining as a standard procedure for ischemic stroke studies(50), we saw significantly smaller infarct volumes in the Ly6G-treated group, consistent with functional scores and microcirculatory parameters (Fig 5/g-h).

Finally, in another group of mice, we wanted to evaluate the histological damage at periinfarct zones at a microscopic scale. Following 1 hour-transient dMCAO, we immediately treated the animals with either anti-Ly6G (n=3) or isotype control antibody (n=3), this time without opening a frontoparietal cranial window to eliminate any possible direct effect of surgery on the vasculature. The next day, they were injected with intraperitoneal propidium iodide (PI), four hours before they were transcardially perfused with a fluorescent gel containing fluorescein isothiocyanate (FITC)-albumin conjugate, at the 24^th^ hour after reperfusion. PI is a dye that is normally cell impermeable but can pass into the cells if membrane permeability is increased in the case of cellular damage, allowing the dye to interact with intracellular nucleic acids and become fluorescent. This approach was used previously to stain degenerating neurons in an *in vivo* cerebral ischemia model(51). Fluorescent gel perfusion and optical index matching with fructose solutions allowed high-resolution ex vivo imaging of the microvasculature (52, 53) (Fig 5/i-k) while retaining the three dimensional structure of the brain. We were able to map the vasculature to identify the MCA-supplied region and identify the injured cells with their PI fluorescence. No specific PI staining was observed in nonischemic zones of the ipsilateral hemisphere (Fig 5/i). We counted the PI-positive cells at the periinfarct zone within 500 µm distance to the nonischemic zone, at two different ROIs in each animal. We determined the number of healthy nuclei taking advantage of their shadows in the FITC-channel over the background signal (Fig 5/i). We did not distinguish between neuronal and other cellular nuclei for this analysis. We found a significant reduction in the fraction of degenerating cells in this zone of potential for recovery with treatment, in the Ly6G treated group (Fig 5/i-j), as a histological hallmark of the observed effect in microcirculation and functional scores.

## Discussion

Using direct in vivo imaging strategies for longitudinal monitoring of microcirculation and tissue injury, we present a novel dynamic mechanism for persistent dysfunctional microcirculation in tissues that can be potentially rescued, and we attributed a role to a hyperacute neutrophil involvement in this dysfunction by specific antibody-mediated targeting of the neutrophils. Technically, evaluation of dynamic capillary stalls was only possible by the high spatiotemporal resolution imaging of OCT angiogram time-series, which clearly indicated flowing RBC signal within hundreds of capillaries simultaneously without any need for labeling.

The experimental power of this study partly originates from the well-controlled and carefully monitored occlusion and recanalization of dMCA along with detailed classification of the ischemic tissue and analyzing the tissue with identical severity of ischemic involvement across groups. The way we distinguish between irreversibly damaged and salvageable tissues was not based simply on blood flow changes but on an integrated approach considering capillary perfusion level and in vivo optical changes in the tissue with histological validation. The ability to map the signal attenuation across the cortex allowed in vivo demonstration of irreversibly damaged tissue, in line with previous reports on the histologically validated abnormal scattering OCT signal attenuation profile (39, 84). While the exact histopathological hallmark of this optical phenomenon is reserved for future investigation, it was extremely useful for evaluation of the extent of ischemic damage without sacrificing the subject and allowed us to adjust for the natural variability in experimental stroke because of the differences in vascular anatomy and collateral supply. Moreover, unlike other methods of transient dMCAO (like thromboembolic, microclip, suture-ligation, electrocoagulation(54–57)), we could precisely achieve full recanalization at every attempt with minimal variability. The experimental model therefore can also be considered relevant for endovascular stroke treatment perspective.

By expanding our previous observations under physiological conditions(17), we see that the capillary RBC flow is extremely irregular and intermittent in the ischemic penumbra, with cells repeatedly getting stuck and released in the microcirculation. We already know that within brain tissue, capillary flow velocities are very different from each other, plasma and red blood cells are differentially partitioned at bifurcations(58), causing a spatial heterogeneity which can lower oxygen extraction efficiency(16, 59–62). Simulations have revealed that the flow kinetics and partitioning in the tight lumen of capillaries are dominated by complex physical cell-cell interactions and rheological parameters which are influenced by temporal events affecting the incidental cell distributions(58). Our findings introduce experimental data on the temporal heterogeneity in individual capillary segments, which can readily influence the upstream and downstream vessels in the network, and cumulatively in a chain-like fashion. As a slight change in stall dynamics can have a profound impact on the overall microvascular blood flow(63) and oxygenation, this can be considered analogous to a traffic jam formation initiated by slight vehicle movement disturbances in a road network(19, 20). We show that these dynamic disturbances are not simple by-standers but can be responsible for ongoing tissue damage in potentially recoverable tissues under acute ischemic conditions. This is in line with a recent study showing abnormal vasospasms in hippocampal capillaries during seizure-related oxidative stress associated with nearby neurodegeneration(64). While single rare capillary stalls, as in healthy animals(17) could possibly be tolerated by the capillary network to some extent, in high numbers and under oligemic conditions with decreased functional capillary reserve, they can aggravate tissue injury in the initial hours of ischemia. Most importantly, this dysfunction is very profound in the penumbra after recanalization, which can help explain the extended neurological impairment or less-than-expected clinical benefit(6, 7). Our data may bring an insight on how the ischemic penumbra can suffer a considerable amount of damage, possibly contributing to the previously shown selective neuronal necrosis accumulating over days after reperfusion(65–67). The extent of this penumbral injury can cause further functional impairment, by delaying recovery, or lowering cognitive reserve(66).

The phenomenon of capillary plugs in ischemia by red and white blood cells is not entirely new. Progressively increasing leukocyte plugging has already been shown in skeletal and myocardial muscle after ischemia and reperfusion, increasing microvascular resistance, negatively affecting the blood flow(68–70). Neutrophils have previously been shown to increase their endothelial adhesion after stroke(38, 71) and occlude the capillary bed(72). Endothelial adhesion molecules like VCAM1 and P-selectin are rapidly upregulated under ischemic conditions as a possible trigger for increased leukocyte-endothelium interactions(38, 73–75). Neutrophils can extend tissue injury by permanently occluding capillaries and also by releasing proteases after their recruitment and passage into the brain parenchyma (14, 15, 72, 76, 77). What we are proposing here is the functional significance of varying levels of flow heterogeneity introduced into the microcirculation at least by neutrophils. Nevertheless, improving the temporary interruptions in capillary flow in the already (but suboptimally) flowing capillaries may be a more compelling target to positively influence the overall cerebral perfusion and oxygen availability, rather than reopening a considerable portion of permanently occluded capillaries to improve the microcirculation. For now, we cannot exclude the contribution of other factors like interactions between platelets, RBCs and other leukocyte types. The detailed principles governing the ultrastructural changes in the penumbral tissue causing an extremely high amount of permanent capillary occlusions and temporary stalls would be an exciting topic for further investigation.

Anti-Ly6G antibodies are traditionally used for in vivo depletion of neutrophils in mice, an effect usually established within 24 hours(47, 49). We decided to use the neutrophil-targeting approach based on our in vivo TPM observations that leukocytes were dynamically plugging the ischemic capillaries in penumbra getting stuck in ischemic microcirculations (Supplementary Movie 3, 4 and 5), to see if we could improve capillary flow patterns along with tissue survival, in line with the recent observations on a mouse model with Alzheimer’s disease(87). Modification of dynamic capillary stalls was indeed possible even with this treatment being applied after recanalization, one hour after ischemia onset and without the requirement for absolute reduction of neutrophil count. The application of monoclonal antibody immediately after recanalization is particularly relevant for clinical cases, as it modelled an adjuvant therapy to mechanical thrombectomy.

This is not the first study targeting neutrophils in ischemic stroke and other models of neurological diseases. Most of the previously published studies focused on decreasing the neutrophil counts or limiting recruitment, to prevent tissue damage by their contribution to BBB leakage, reactive oxygen species and toxic enzyme release within the parenchyma(88). These studies focused mostly on an effect happening over a few days after ischemic induction. Typically recruitment into brain parenchyma begins 6-8 hours after ischemia(88, 89) and peaks at 24-48 hours(76, 88, 90). Nevertheless, the majority of these studies reported a beneficial outcome in animal models(88). If we consider the anti-Ly6G that was used in this study, targeting of neutrophils by the same antibody was found beneficial for subarachnoid hemorrhage-related vasospasm in one study(91) and also improved early parenchymal blood flow in another(92). The same antibody was also found to be improving functional scores after stroke at 72 hours in hyperlipidemic mice(93). Our results, despite confirming an effect by neutrophil targeting, should be interpreted differently as they indicate a rather hyper acute direct effect on microcirculatory flow patterns with early administration of the anti-Ly6G monoclonal antibody simultaneously with full recanalization.

One potential limitation of our study was the reliance on acute surgical preparations and the necessity to use an inhalational anesthetic, isoflurane for our long-duration experiments. Although we utilized our best efforts to control for the confounders, monitored systemic physiological parameters, and included proper sham and control groups for appropriate comparisons, we cannot completely exclude any effect of anesthesia or acute effects of surgical trauma on our observations. Isoflurane, for instance, is a neuroprotectant and has profound effects on cerebral blood flow(94–98). The acute surgical procedures that involved craniotomy may have aggravated inflammatory changes in the tissue, increasing leukocyte adhesion(71). These issues will need to be addressed in future studies, that would be conducted in animals with chronic cranial windows and ideally in the awake setting. Another limitation was inclusion of relatively young animals to limit the variability related to the factors in the care of animals that can occur throughout lifetime. Since most stroke cases occur in elderly patients with other comorbidities, the extension of this study’s findings in such mice with appropriate cardiovascular disease models will be necessary. Although we are well aware of the benefit of both male and female mice for experimental studies, we had to include only male mice here, because the well-documented gender effect on neutrophil function as well as the high variability of sex hormones in female mice(21), that would increase the number of mice to be used several folds. From our perspective, this was acceptable as we were not proposing an established treatment for immediate clinical translation, but we were rather defining a dynamic pathogenic mechanism in a rather homogeneous experimental sample. Finally, and maybe the most importantly, our therapeutic target, the Ly6G protein, is expressed only in mice and not in humans(99). Therefore, this molecular target is not directly applicable for human use. Since Ly6G’s endogenous ligand and exact actions are still unknown, it will take time before we identify a working candidate for human use. Deciphering the exact mechanism of action of this antibody was beyond the scope of this research but proposes an interesting research question for investigators in this field. At least for now, the strong association of neutrophil/lymphocyte ratio in blood with severity of stroke in stroke patients is encouraging for our experimental observations (100–103).

It can be argued that our observed effect on capillary flow was generated by induced leukopenia, with less cells in circulation that can potentially occlude the capillary flow. However, we noticed a significant effect as early as 2 hours after recanalization, even before settlement of full-blown neutropenia. This implies that absolute reduction of neutrophil counts may not be necessary for the effect. We are cautious about making interpretations about any effect of Ly6G antibodies on neutrophil adhesion, size or flexibility, before they are fully depleted, as we did not have the capabilities for a systematic and valid investigation in this field. Nevertheless, Ly6G targeting was previously suggested to decrease the endothelial adhesion of leukocytes, as previously suggested by a mechanism over downregulation of B2-integrins on the neutrophil surface(104).

These findings, in our opinion, are very helpful for better understanding the dynamic flow issues at capillary level in cerebral ischemia. It may be useful to imagine those brief but frequent and persistent ‘traffic jams’ while interpreting any structural change in microcirculation or any change in hematological parameters. This improvement in the field may lead to novel strategies to improve overall cellular passage through the microcirculation to enhance the clinical benefits of endovascular treatment. Possible targets may not be just limited to neutrophil count and adhesion and would also include deformability of blood cells, endothelial surface protein expression and reactive oxygen species. Our findings may inform further studies on the microcirculation disturbances for other acute brain disorders, like migraine, epilepsy and concussion. We believe that recognition of this dynamic phenomenon widely in the medical community would catalyze more basic and clinical research studies and further development in the near future.

## Materials and Methods

### Animals and Surgical Procedures

All experiments were approved by the Institutional Animal Care and Use Committees of Massachusetts General Hospital and Boston University. Experiments were conducted following the Guide for the Care and Use of Laboratory Animals.

For this study, 12-16-week old C57BL/6 mice (24-30g, male, purchased from Charles River Laboratories) were used. Animals were housed under diurnal lighting conditions with free access to food and water. Surgery and experiments were performed under isoflurane anesthesia (2-3% induction, 1-2% maintenance, in 60% Nitrogen, 40% Oxygen mixture). The left femoral artery was cannulated for blood pressure measurements, except for the animals intended survive overnight. Body temperature was maintained with a homeothermic blanket control unit (Harvard Apparatus). Following a midline skin incision, and removal of scalp and periosteum, a custom-made aluminum bar was glued over the left half of the skull, for fixation of the head. Then,, the temporalis muscle was dissected to reveal the squamous portion of the temporal bone. The temporal bone over the distal MCA, 1 mm above the zygomatic arch was drilled and a craniotomy of 2 mm diameter was opened, with dura intact. Next, another craniotomy was opened (4 mm in diameter) over the frontoparietal area (center: approximately 3 mm lateral and 1 mm posterior to bregma), over the edge of the MCA-supplied cortex, including the MCA-ACA border zone region. The cortex was then covered with 0.7% agarose solution in aCSF, followed by a 5-mm glass coverslip. The window was sealed with dental cement. The animal was then placed under the imaging system. We waited for at least an hour for stabilization of the cerebral blood flow until we started the baseline imaging.

Mean arterial pressure was monitored in animals with femoral cannulation. Arterial oxygen saturation and heart rate was monitored noninvasively with a paw probe (MouseSTAT Jr., Kent Scientific Instruments). We did not make attempts to monitor arterial blood gases or end-tidal pCO_2_ levels. Because, firstly, the animals were spontaneously breathing during the experiments and therefore close monitoring of these parameters for fine adjustments like ventilator settings was not necessary. In fact, excessive invasive manipulations like tracheostomy and positive pressure ventilation as well as a decrease in blood volume as a result of repeated arterial blood samplings would increase experimental variability and also generate questions on whether the observed findings were pathophysiological or simply methodologically introduced. Our consistent observation that every animal to survive overnight recovered from isoflurane anesthesia within 60 seconds of discontinuation of gas flow was accepted as an indicator against any respiratory acidosis due to the experiments.

At the end of the experiments (2 or 24 hours after reperfusion) animals were decapitated and the brains were removed quickly for histological confirmation of the infarct through TTC staining (see below). In a separate group (n=3 per treatment group), animals were transcardially perfused with a fluorescent gel containing fluorescein isothiocyanate (FITC)-albumin conjugate (composition in detail below), 4 hours after intraperitoneal injection of propidium iodide (300µl of 1 mg/ml PI solution [Molecular Probes] diluted with 300 µL injectable sterile saline). The animal was then decapitated and the brain was extracted for further processing (see below).

### Transient Distal MCAO Model

For initiation of ischemia, an occlusion device (Fig 1) made by the fusion of two blunted borosilicate glass micropipettes was be positioned over the distal MCA in the recently opened temporal craniotomy, with help of a micromanipulator (YOU-1, Narishige, Japan)(23). After baseline imaging of the chronic cranial window was complete, the pipette was gently advanced to press on the MCA trunk, at the point where it crossed the inferior cerebral vein, until blood flow dropped. Cerebral blood flow was monitored via laser speckle blood flowmetry in real-time to confirm ischemia initiation and maintenance. After one hour of ischemia, the pipette was drawn back to recanalize the artery. In sham animals, all surgical procedures were performed, but the blunted pipette compression was applied over the dura adjacent to the MCA, with no compression of arterial or venous structures.

### Antibody Treatment

For targeting of leukocytes, anti-Ly6G antibody (BD Biosciences) 2mg/kg was administered through the retroorbital sinus over 30 seconds, ∼3 min after recanalization. For the control group, isotype control antibody, 2mg/kg (BD Biosciences) was injected. 100 µL of antibody solution was diluted with 50 µL of sterile injectable saline before injection. Injection was performed by the investigator performing the dMCAO procedure who was blinded to the injection type.

### Optical Coherence Tomography

A spectral-domain OCT system (1310 nm center wavelength, bandwidth 170 nm, Thorlabs Telesto III) was used for imaging of the cerebral cortex. OCT-angiograms were constructed by a decorrelation-based method(27). In this study, imaging regions of interest (ROI) were raster scanned at different pixel resolutions. Maximum intensity projections (MIP) over a depth range of 150-250 μm beneath the brain surface were extracted from angiogram volumetric data for analysis as these layers provided a high-quality angiogram signal from capillaries. When RBCs stall in a capillary segment, the draining segment disappears from the OCT-angiogram as no dynamic signal is detected. The disappearance and reappearance of individual segments could be readily identified as a stalling segment/event. We marked and manually counted each stall for quantification (see below). Phase resolved Doppler OCT (prD-OCT)(28) was utilized to quantify the axial blood flow velocity in penetrating vessels. Please refer to the Supplementary Online Material for details on OCT acquisition and data processing.

For each timepoint during the experiments (baseline, ischemia, reperfusion-at 1 hour, 2 hours and 24 hours) a new OCT dataset was acquired. Each OCT dataset consisted of one time series (60 consecutive volumes) of narrow-field data, 5 volumes of wide-field data (both for 5x and 10x) and one volume of middle-field PrD-OCT data. The same cortical region was repeatedly imaged for experimental timepoints and readjustments were not necessary since the animal was not removed from the imaging system during experiments (apart from repeat imaging at 24 hours after reperfusion). On a few occasions minor shifts in ROI location was necessary after baseline acquisition, to acquire the targeted zone of the core-penumbra boundary after onset of ischemia. The imaging field was not changed once ischemia was initiated.

### Two Photon Microscopy

A commercial laser scanning TPM system (Bruker Investigator) with an integrated Becker and Hickl TCSPC FLIM module was used with a Nikon, 16X, 0.8NA, water-immersion objective. During experiments, the objective was heated using a TC-1-100s objective heater from Bioscience Tools to keep the immersion water temperature constant around 37°C. Oxygen-sensitive dye PtP-C343 (60 mg/ml, 120 µL) was retro-orbitally injected ∼30 min before imaging. Capillary fluorescence was initially imaged for anatomic guidance. Time-series of capillary flow to demonstrate the stalls in supplementary videos 3-4 were acquired with 4.0 optical zoom, 512×512 pixel resolution over ∼200×200 µm2 field of view. The image frame interval was 1 second. PtP-C343 fluorescence visualized the plasma, while blood cells remained unstained. For in vivo visualization of leukocytes, 100 µL of Rhodamine-6G (0.05% in PBS, freshly prepared) was injected retro-orbitally, 5 min before imaging. The dye was excited at 830 nm. Five adjacent regions at 150 µm depth in the penumbra next to the infarct core were imaged for 200 seconds in each animal, at 1 frame per second. Each region was again ∼200×200µm^2^, 512×512 pixels. Please refer to the Supplementary Online Material for details on fluorescence and phosphorescence data acquisition.

pO_2_ determination was done at all visible microvasculature segments, including precapillary arterioles, capillaries and postcapillary venules, across a single focal plane (at 120-170 µm deep) over an ∼440×440 µm imaging field of view. 20-30 points inside capillaries were measured at each imaging region, the exact number depending on the number of visible capillaries. For each animal, four separate regions were imaged and the measured data were pooled together. At each point location inside the vasculature, we summed the phosphorescence decays of 3000 trials, which were then averaged across 3 repetitions. During postprocessing, phosphorescence lifetimes (taus, τ) were determined by fitting to an exponential decay model, which was then converted to pO2 values via calibration plots for the PtP-C343 dye(32). For time-series pO2 measurements in selected capillaries, we longitudinally imaged the same points with no repeats with a ∼1s temporal resolution, for ∼150 seconds in total. Three consecutive measurements were averaged together (covering ∼3 seconds cumulatively) to correct nonspecific fluctuations of measured pO_2_ values. Ex vivo brain samples were mounted in a petri dish inside the index-matching fructose solution. With the abovementioned excitation and emission parameters, Z-stacks of ∼400×400×300 µm^3^ with 3 µm slicing interval were acquired in the periinfarct areas corresponding to the zones investigated with OCT. Images from both red fluorescence (for PI) and green fluorescence (for FITC) were simultaneously acquired. In each animal, two nonoverlapping regions were imaged.

### Quantification of Stalls

Quantitative analyses of capillary stalls were performed as reported previously(17). In angiogram time-series, stalling capillary segments could be easily identified as a sudden drop in signal intensity; disappearance and reappearance of flowing RBCs in individual segments could be observed. We marked each stall for quantification using custom code in MATLAB. This software plotted the signal intensity fluctuations in a segment identified to be stalling, making detection of individual stall events easier. Stall incidence was calculated by the number of segments stalling at any time in the time-series divided by the total number capillaries. Stall point prevalence indicated the average number of stalled segments in each imaging frame divided by the total number of capillaries. Cumulative stall duration was expressed as the percentage of time that a given stalling capillary segment was not flowing.

### Statistical Analyses

All manual data analyses were performed by a blinded fashion with respect to the experimental group (isotype-control or anti-Ly6G antibody injected). Independent groups (isotype control vs anti-Ly6G) were compared by Mann-Whitney U test unless otherwise indicated. Dependent comparisons between different time points were done by Friedman test followed by Dunn’s multiple comparisons test for comparisons with baseline. P<0.05 was accepted as statistically significant. P values between 0.05 and 0.09 were reported as near-significant. Results were expressed as mean ± standard error of the mean (SEM), unless otherwise indicated.

## Acknowledgements

This study was supported by National Institute of Health grants R01-EB021018 and R01-NS108472. Şefik Evren Erdener’s work was additionally supported by the Turkish Neurological Society and Hacettepe University Scientific Research Projects Coordination Unit (TUI-2019-18106). Dmitry Postnov was supported by the grant from Novo Nordisk Foundation, Denmark (NNF17OC0025224). We would like to tank Andrew K. Dunn (University of Texas, Austin) for laser speckle acquisition software, Sergei Vinogradov (University of Pennsylvania) for providing oxygen-sensitive phosphorescent probes and for Michael A. Moskowitz (Massachusetts General Hospital) for helpful discussion and comments. We also would like to thank Blaire S. Lee for her technical assistance.

