## Supplementary Material for "Neutrophil-mediated dynamic capillary stalls in ischemic penumbra: persistent traffic jams after reperfusion contribute to injury"

**SUPPLEMENTARY ONLINE MATERIAL**

**SUPPLEMENTARY METHODS**

*Animals and Surgical Procedures*

All experiments were approved by the Institutional Animal Care and Use Committees of Massachusetts General Hospital and Boston University. Experiments were conducted following the Guide for the Care and Use of Laboratory Animals.

For this study, 12-16-week old C57BL/6 mice (24-30g, male, purchased from Charles River Laboratories) were used. Animals were housed under diurnal lighting conditions with free access to food and water. Surgery and experiments were performed under isoflurane anesthesia (2-3% induction, 1-2% maintenance, in 60% Nitrogen, 40% Oxygen mixture). The left femoral artery was cannulated for blood pressure measurements, except for the animals intended survive overnight. Body temperature was maintained with a homeothermic blanket control unit (Harvard Apparatus). Following a midline skin incision, and removal of scalp and periosteum, a custom-made aluminum bar was glued over the left half of the skull, for fixation of the head. Then, approaching between the right lateral epicanthus and external auditory meatus, the temporalis muscle was dissected to reveal the squamous portion of the temporal bone. The temporal bone over the distal MCA, 1 mm above the zygomatic arch was drilled and a craniotomy of 2 mm diameter was opened, with dura intact. To prevent overheating, extended drilling of the same area for more than 2-3 seconds was avoided and the skull was cooled with aCSF at room temperature throughout the drilling process. The exposed MCA and surrounding cortex were flushed with artificial CSF (aCSF) and then was covered with aCSF-soaked-gauze for protection. Next, another craniotomy was opened (4 mm in diameter) over the frontoparietal area (center: approximately 3 mm lateral and 1 mm posterior to bregma), over the edge of the MCA-supplied cortex, including the MCA-ACA border zone region was opened. The cortex was then covered with 0.7% agarose solution in aCSF, followed by a 5-mm glass coverslip. The window was sealed with dental cement. The animal was then placed under the imaging system. We waited for at least an hour for stabilization of the cerebral blood flow until we started the baseline imaging.

During the entire length of the surgical procedure, the animal was heated by a homeothermic blanket with rectal-probe feedback to maintain temperature at 37 °C. Mean arterial pressure was monitored in animals with femoral cannulation. Arterial oxygen saturation and heart rate was monitored noninvasively with a paw probe (MouseSTAT Jr., Kent Scientific Instruments). We did not make attempts to monitor arterial blood gases or end-tidal pCO_2_ levels. Because, firstly, the animals were spontaneously breathing during the experiments and therefore close monitoring of these parameters for fine adjustments like ventilator settings was not necessary. In fact, excessive invasive manipulations like tracheostomy and positive pressure ventilation as well as a decrease in blood volume as a result of repeated arterial blood samplings would increase experimental variability and also generate questions on whether the observed findings were pathophysiological or simply methodologically introduced. Our consistent observation that every animal to survive overnight recovered from isoflurane anesthesia within 60 seconds of discontinuation of gas flow was accepted as an indicator against any respiratory acidosis due to the experiments.

At the end of the experiments (2 or 24 hours after reperfusion) animals were decapitated and the brains were removed quickly for histological confirmation of the infarct through TTC staining (see below). In a separate group (n=3 per treatment group), animals were perfused with a fluorescent gel containing fluorescein isothiocyanate (FITC)-albumin conjugate (composition in detail below), 4 hours after intraperitoneal injection of propidium iodide (300µl of 1 mg/ml PI solution [Molecular Probes] diluted with 300 µL injectable sterile saline). For this perfusion procedure, under isoflurane anesthesia the abdominal wall and peritoneum were dissected, the incision was extended to the rib cage on the lateral edges and the diaphragm was cut open. The sternum was retracted to expose the chest cavity and the heart. The left ventricle was then pierced with a shielded-winged needle (21G). The right atrium was cut with spring scissors to allow drainage of blood. Then, 30 ml of 20 u/mL heparin sodium in phosphate-buffered saline (PBS) was injected at a rate of 30 mL/min. Next, 15 mL of a fluorescent perfusion gel (containing 10% gelatin, 0.4% FITC-albumin ) at 42 °C was injected into the left ventricle, similar to a previously reported protocol(22). The animal was then immersed in ice for 10 min in a head-down position to keep the solidifying gel inside the cranial vasculature. The animal was then decapitated and the brain was extracted for further processing (see below).

*Transient Distal MCAO Model*

For initiation of ischemia, an occlusion device (Fig 1) made by the fusion of two blunted borosilicate glass micropipettes was be positioned over the distal MCA in the recently opened temporal craniotomy, with help of a micromanipulator (YOU-1, Narishige, Japan)(23). Micropipette was positioned so that it was exactly perpendicular to the brain surface plane. The line connecting the tips of the occlusion device was making a 30° oblique angle with the longitudinal axis of MCA. After baseline imaging of the chronic cranial window was complete, the pipette was gently advanced to press on the MCA trunk, at the point where it crossed the inferior cerebral vein, until blood flow dropped. Cerebral blood flow was monitored via laser speckle blood flowmetry in real-time to confirm ischemia initiation and maintenance. After one hour of ischemia, the pipette was drawn back to recanalize the artery. In sham animals, all surgical procedures were performed, but the blunted pipette compression was applied over the dura adjacent to the MCA, with no compression of arterial or venous structures.

*Antibody Treatment*

For targeting of leukocytes, anti-Ly6G antibody (BD Biosciences) 2mg/kg was administered through the retroorbital sinus over 30 seconds, ~3 min after recanalization. For the control group, isotype control antibody, 2mg/kg (BD Biosciences) was injected. 100 µL of antibody solution was diluted with 50 µL of sterile injectable saline before injection. Injection was performed by the investigator performing the dMCAO procedure who was blinded to the injection type.

*Laser Speckle Contrast Imaging*

Laser speckle contrast imaging (LSCI) was performed as described previously(24) with minor alterations. Briefly, the right-sided cranial window was illuminated using volume holographic grating (VHG) stabilized laser diode (785nm, LP785-SAV50, Thorlabs)(25) with a power density of approximately 20mW/cm^2^ (26). Backscattered light was recorded using a 10X (~2.8x2.8 mm area) and a 5X objective (~5x5 mm area), with a 1600x1600 subset of pixels of a CMOS camera (Basler acA2040-180kmNIR, 2048x2048 pixels, 4.4 µm pixel size) connected to the OCT imaging microscope with a dichroic mirror. This setup allowed us to acquire OCT and laser speckle data simultaneously. A polarizer was placed after the dichroic to ensure it has no effect on the OCT signal. Camera exposure time was set to 5 ms.  The speckle size was adjusted by altering the pupil diameter of an iris in the detection path to achieve the optimal signal to noise ratio. Fifteen raw images were acquired every 1 second, converted to speckle contrast (K) images and averaged online using custom software (kindly provided by Andrew Dunn, University of Texas in Austin).

*Optical Coherence Tomography*

A spectral-domain OCT system (1310 nm center wavelength, bandwidth 170 nm, Thorlabs Telesto III) was used for imaging of the cerebral cortex. Axial resolution of the system in air was 4.6 μm. Given the light refractive index of 1.35 for brain tissue, the axial resolution in brain was ~3.5 μm. The imaging speed was 76,000 A-scan/s. A 10X objective (10X Mitutoyo Plan Apochromat Objective, 0.28 NA) and a 5X objective (5X Mitutoyo Plan Apochromat Objective, 0.14 NA) were used in this study allowing a transverse resolution of 3.3 μm and 7 μm, respectively.

OCT-angiograms were constructed by a decorrelation-based method(27). While conventional structural OCT imaging acquires one *xz* B-scan for each *y* position, the decorrelation-based method repeats two B-scans and then analyzes the differences in the image intensity and phase between the repeated B-scans. There would be no difference for repeated voxels for static tissue. In contrast, signal from dynamic tissue, such as a blood vessel, will fluctuate between repeated B-scans due to particle movement (e.g. flowing RBCs). The subtraction between two consecutive B-scans will make the dynamic blood vessels stand out as bright regions in the OCT-angiogram. A 2D raster scan in the X-Y plane will lead to a 3D OCT-angiogram which presents the microvascular structure. In this study, imaging regions of interest (ROI) were raster scanned at different pixel resolutions. For stall time-series, data were acquired with a 400$\times$400-pixel count in X-Y plane (covering a 600$\times$600$\times500$ ${\mu m}^{3}$ ROI, using 10X objective; narrow ROI) for stall time series. For wide-field angiograms, acquisition was done with a 500$\times$500 pixel count (2000$\times20$00$\times500$ ${\mu m}^{3}$using 10X objective and 2500$\times25$00$\times500$ ${\mu m}^{3}$ using 5X objective; wide ROI). Narrow and wide ROIs were concentrically placed, with the same center coordinates. Each 400$\times$400 pixel OCT-angiogram acquisition took ~6.5 seconds, and a time-series of 60 volumes of the cortical microvasculature was consecutively acquired for each ROI, this acquisition lasted ~6.5 min. 500$\times$500 pixel-resolution angiograms took ~9 seconds. Maximum intensity projections (MIP) over a depth range of 150-250 μm beneath the brain surface were extracted from angiogram volumetric data for analysis as these layers provided a high-quality angiogram signal from capillaries. When RBCs stall in a capillary segment, the draining segment disappears from the OCT-angiogram as no dynamic signal is detected. The disappearance and reappearance of individual segments could be readily identified as a stalling segment/event. We marked and manually counted each stall for quantification (see below).

Phase resolved Doppler OCT (prD-OCT)(28) was utilized to quantify the axial blood flow velocity of arterials and venous. prD-OCT data was acquired using an M-mode scanning pattern (i.e. repeat A-scans for each (*x,y*) position), and was performed in the same ROI as the OCT-angiogram. We performed 100 repeated A-scans for each transverse position (X, Y) to extract blood flow velocity. The imaging field was scanned with 400$\times$400 (X, Y) transverse positions, and the ROI spanned $1000\times1000\times500$ ${\mu m}^{3}$ (middle ROI). The center of this ROI was identical with the narrow and wide ROIs used for angiogram imaging. This pixel count and imaging field dimensions optimized the number of visible penetrating arterioles (as more arterioles identified would ensure more accurate estimation of cerebral blood flow in this region) while keeping data size and acquisition time reasonable. The acquisition time for 100 A-scans was ~1.3 ms which was short enough to avoid motion artifacts primarily resulting from cardiac pulsation (71-194 ms(29))and respiration (261-750 ms(30)). The total time to acquire one volume of prD-OCT data was ~8 min. Offline data processing for prD-OCT was performed with MATLAB on a 16,100-core cluster system provided by Boston University. Data spanning 20-250 $\mu m$ beneath the brain surface was extracted from each A-line. Since the flow of RBCs causes the OCT signal phase change and it is opposite in penetrating and ascending vessels, we use such phase change information to identify arteries/arterioles from veins/venules. Axial blood flow velocity was obtained by calculating the weighted histogram of the unwrapped OCT signal phase slop. This method provides robust axial blood flow measurement in both large vessels and capillaries. For details please refer to Tang *et al*(31).

For calculation of cerebral blood flow, a transverse plane passing through the penetrating arterioles was selected (around 50-70 µm below cortical surface). Based on the flow direction, arterioles were first identified and average axial velocity was calculated for each arteriole. Then, blood flow for each arteriole was calculated by taking the product of axial blood flow velocity and the cross-sectional area of the arteriole. Blood flow values calculated from all identified arterioles in the $1000\times1000 {\mu m}^{2}$ area were summed to get the cerebral blood flow. The same penetrating arterioles at the same depth were used for repeated blood flow data analysis acquired at different experimental time points.

To determine the OCT signal attenuation changes, we used a 5X objective to scan 500$\times$500 transverse positions which provided a 3D volume of 2.5$\times$2.5$\times$1 ${mm}^{3}$. The 5X objective had a lower numerical aperture and longer focal depth hence providing more uniform illumination along the depth enabling us to more directly estimate the slope of the exponential attenuation of light with depth. The focus was positioned at the brain surface. To improve the signal-to-noise ratio, four repeated scans were performed to get an averaged OCT signal. We performed a linear fit of the logarithmic OCT axial signal profile between a depth of 50 and 550 µm and calculated the attenuation coefficient as the slope of the fit. Slope values were calculated at each xy position. More negative values indicated a faster axial decay in signal, hence higher scattering, which was characteristic in the well-demarcated infarct tissue.

For each timepoint during the experiments (baseline, ischemia, reperfusion-at 1 hour, 2 hours and 24 hours) a new OCT dataset was acquired. Each OCT dataset consisted of one time series (60 consecutive volumes) of narrow-field data, 5 volumes of wide-field data (both for 5x and 10x) and one volume of middle-field PrD-OCT data. The same cortical region was repeatedly imaged for experimental timepoints and readjustments were not necessary since the animal was not removed from the imaging system during experiments (apart from repeat imaging at 24 hours after reperfusion). On a few occasions minor shifts in ROI location was necessary after baseline acquisition, to acquire the targeted zone of the core-penumbra boundary after onset of ischemia. The imaging field was not changed once ischemia was initiated.

*Two Photon Microscopy*

A commercial laser scanning TPM system (Bruker Investigator) with an integrated Becker and Hickl TCSPC FLIM module was used with a Nikon, 16X, 0.8NA, water-immersion objective. During experiments, the objective was heated using a TC-1-100s objective heater from Bioscience Tools to keep the immersion water temperature constant around 37°C. Oxygen-sensitive dye PtP-C343 (60 mg/ml, 120 µL) was retro-orbitally injected ~30 min before imaging. Capillary fluorescence was initially imaged for anatomic guidance. The dye was excited using femtosecond pulses from a Spectra Physics Insight X3 Titanium:Sapphire laser with a pulse width of ~100fs, 80MHz repetition rate and the output tuned to 920nm and a ConOptics 350-80-LA EOM was used to control laser power. Raster scanning of the laser excitation foci is achieved using a pair of coupled galvanometers scanning over the field of view at a pixel dwell time of 3.2µs. The collected fluorescence is split using a 565nm long pass filter into two light paths. For the reflected path, a 525nm, 50nm bandpass filter is used to visualize green fluorescence while a 620nm, 60nm band pass filter is used to visualize red fluorescence.

Time-series of capillary flow to demonstrate the stalls in supplementary videos 3-4 were acquired with 4.0 optical zoom, 512x512 pixel resolution over ~200x200 µm2 field of view. The image frame interval was 1 second. PtP-C343 fluorescence visualized the plasma, while blood cells remained unstained. For in vivo visualization of leukocytes, 100 µL of Rhodamine-6G (0.05% in PBS, freshly prepared) was injected retro-orbitally, 5 min before imaging. The dye was excited at 830 nm. Five adjacent regions at 150 µm depth in the penumbra next to the infarct core were imaged for 200 seconds in each animal, at 1 frame per second. Each region was again ~200x200µm^2^, 512x512 pixels.

For capillary oxygen pressure (pO_2_) measurements, we excited PtP-C343 with a 10-μs-long train of femtosecond pulses at 920 nm and record the following phosphorescence over a 290-μs-long window. For better collection efficiency due to the shift in the PtP emission spectrum, we replaced the fluorescence filter in the red channel of our detection pathway with a 710nm, 75nm band pass filter. PO_2_ determination was done at all visible microvasculature segments, including precapillary arterioles, capillaries and postcapillary venules, across a single focal plane (at 120-170 µm deep) over an ~440x440 µm imaging field of view. 20-30 points inside capillaries were measured at each imaging region, the exact number depending on the number of visible capillaries. For each animal, four separate regions were imaged and the measured data were pooled together. At each point location inside the vasculature, we summed the phosphorescence decays of 3000 trials, which were then averaged across 3 repetitions. During postprocessing, phosphorescence lifetimes (taus, τ) were determined by fitting to an exponential decay model, which was then converted to pO2 values via calibration plots for the PtP-C343 dye(32). For time-series pO2 measurements in selected capillaries, we longitudinally imaged the same points with no repeats with a ~1s temporal resolution, for ~150 seconds in total. Three consecutive measurements were averaged together (covering ~3 seconds cumulatively) to correct nonspecific fluctuations of measured pO_2_ values.

Ex vivo brain samples were mounted in a petri dish inside the index-matching fructose solution. With the abovementioned excitation and emission parameters, Z-stacks of ~400x400x300 µm^3^ with 3 µm slicing interval were acquired in the periinfarct areas corresponding to the zones investigated with OCT. Images from both red fluorescence (for PI) and green fluorescence (for FITC) were simultaneously acquired. In each animal, two nonoverlapping regions were imaged.

*Quantification of Stalls*

Quantitative analyses of capillary stalls were performed as reported previously(17). In angiogram time-series, stalling capillary segments could be easily identified as a sudden drop in signal intensity; disappearance and reappearance of flowing RBCs in individual segments could be observed. We marked each stall for quantification using custom code in MATLAB. This software plotted the signal intensity fluctuations in a segment identified to be stalling, making detection of individual stall events easier. Stall incidence was calculated by the number of segments stalling at any time in the time-series divided by the total number capillaries. Stall point prevalence indicated the average number of stalled segments in each imaging frame divided by the total number of capillaries. Cumulative stall duration was expressed as the percentage of time that a given stalling capillary segment was not flowing.

*Infarct Size Measurements*

Infarct sizes were quantified by TTC (2,3,4-triphenyltetrazolium chloride) staining. After the animals are decapitated, the brains were quickly extracted and were immersed in ice-cold saline for 2 minutes and then cut into 2-mm-thick coronal slices using a slicing matrix. Sections were incubated in TTC solution (2% in PBS, Sigma) for 30 minutes at 37 C. The infarcts were most prominently visible on the 2^nd^ and 3^rd^ sections, occasionally extending to the 1^st^ section. The slices were immediately imaged with an operating microscope (Nikon SMZ 645, with LabCAM adapter), at the posterior aspect of each section. Infarct zones could be easily identified with lack of TTC staining. In each slice, infarcted cortical area was measured indirectly, by subtraction of the noninfarcted area of the ipsilateral hemisphere from the area of the contralateral hemisphere, to account for any edematous changes in the infarct tissue. Final infarct volume was calculated by multiplying the sum of infarct areas in each slice with 2 (since section thickness was 2 mm).

*Leukocyte Counts*

Blood was withdrawn from the femoral artery catheter for non-survival or from the tail vein for the survival experiments. 5µL of blood was mixed with 195 µL of Turk’s solution (2% acetic acid with methylene blue), and then was spread over a hemocytometer (Hausser Scientific). Leukocytes were identified and neutrophils were differentiated with their nuclear morphology. Total leukocytes and neutrophils were manually counted over 5 regions.

*Neurological Scoring*

Garcia neurological scoring was performed 1 hour after recovery of the animals and approximately 22 hours after ischemia onset, as previously described(33).

The adhesive tape removal test(34) was performed with slight modifications. The mouse

was allowed to get accustomed to the testing cage for 5 minutes. Small adhesive tape bands (3x4 mm) were applied to the non-impaired right forepaw of the mouse. Then the mouse was placed into the testing box and the timer was set. Time needed to remove the adhesive tape (in seconds) was recorded in 5 successive trials, in alternating limbs (both affected and unaffected sides), following two initiation trials. The median of these 5 trials were used as the final score for each limb of each animal.

*Ex Vivo Tissue Clearing*

Extracted brains were kept in 4% PFA in PBS overnight, at room temperature, followed by a 5-day incubation in PBS. Then, the brains were subsequently immersed in 20, 40, 60, 80 and 110% fructose solutions in distilled water containing 0.5% α-thioglycerol, with gentle shaking, for 4, 6, 8, 12 and 24 hours, respectively. 110% saturated fructose solution, also known as SeeDB(35) allowed deep tissue penetration for two-photon imaging by optical index matching of the tissue.

For analysis, PI-positive cells were manually counted using the cell counter plug-in of FIJI. Yhe number of unstained cells were determined by taking advantage of the dark appearance of cell bodies over background neuropil fluorescence in FITC images. Those bodies were segmented by inverse thresholding in image stacks they were volumetrically counted by 3dObjects Counter plugin(36) of FIJI.

*Statistical Analyses*

All manual data analyses were performed by a blinded fashion with respect to the experimental group (isotype-control or anti-Ly6G antibody injected). Independent groups (isotype control vs anti-Ly6G) were compared by Mann-Whitney U test unless otherwise indicated. Dependent comparisons between different time points were done by Friedman test followed by Dunn’s multiple comparisons test for comparisons with baseline. P<0.05 was accepted as statistically significant. P values between 0.05 and 0.09 were reported as near-significant. Results were expressed as mean ± standard error of the mean (SEM), unless otherwise indicated.

**SUPPLEMENTARY FIGURES**

**Supplementary Figure 1**

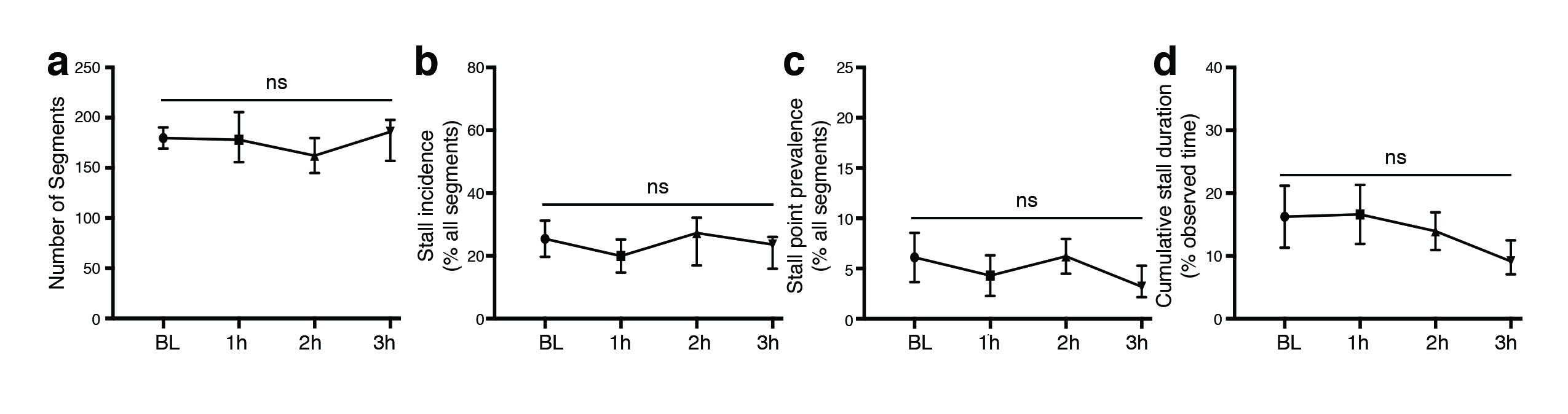

Sham experiments (n=4). Capillary stall parameters, despite showing some variability because of the effect of acute cranial window preparation and anesthesia, did not show a significant change over 3 hours of observation. The trend in cumulative stall duration to decrease over time (d) supports that increased stall duration after ischemia-reperfusion is not due to the surgical preparation. Data expressed as mean ± SEM.

**Supplementary Figure 2**
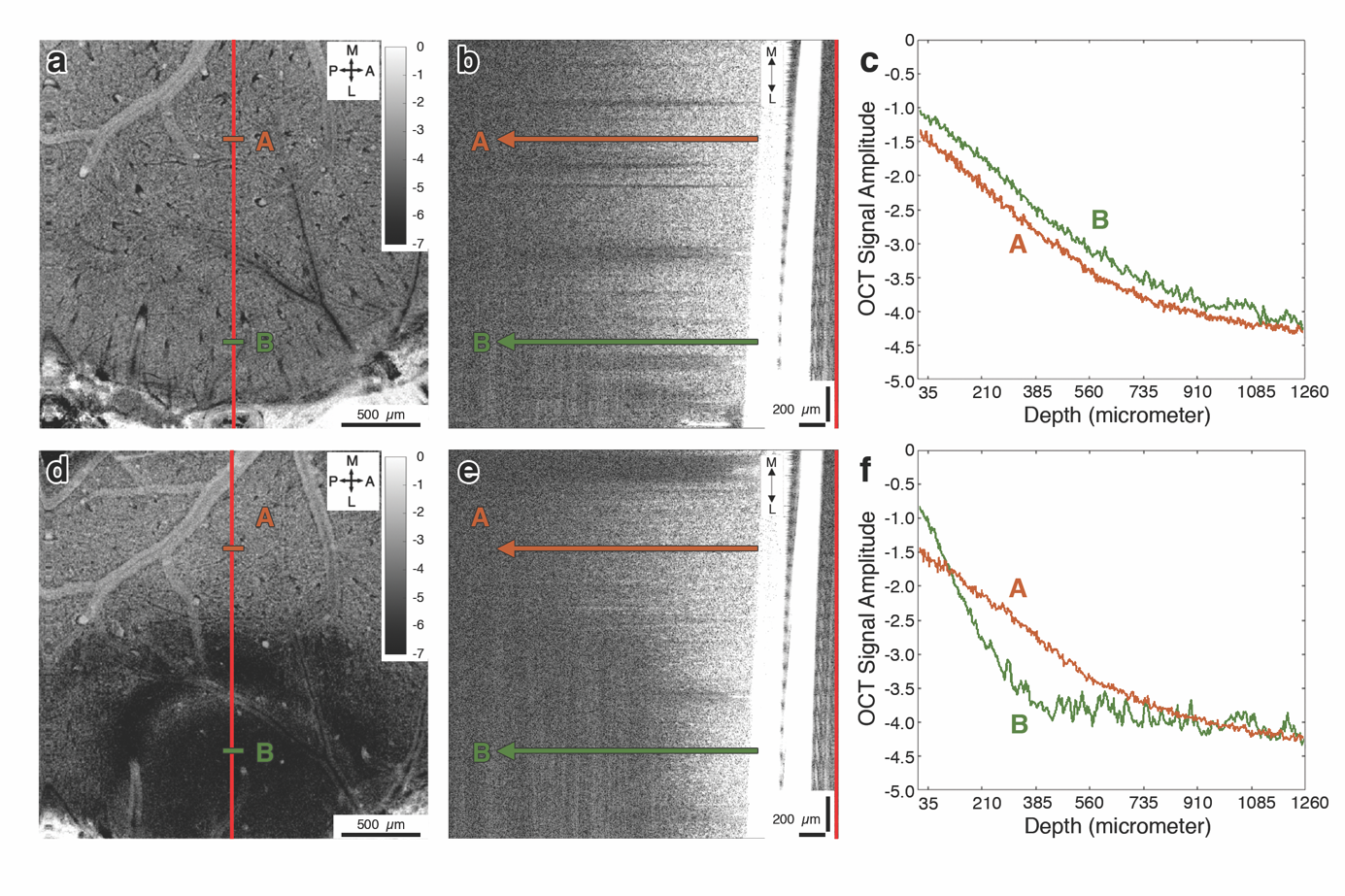

**(a)** Baseline signal attenuation slope map shows uniformity across the cranial window. **(b)** Raw OCT amplitude signal at the level of the red vertical line also confirms the uniform scattering profile. **(c)** Two plots along the A and B lines show similar slope of the depth-dependent signal decay (note that the OCT signal is on a logarithmic scale). **(d-f)** However after establishment of ischemic injury, in the lateral side of the cranial window supplied extensively by the MCA, the core, there is abnormally very low signal penetration which is indicated by a higher decay slope of line-B, compared with that in the penumbra (line-A).

Scale bars in axial views: 500 µm, in coronal views: 200 µm. Unit of calibration bars in (a) and (d) is 1/mm^2^.

**Supplementary Figure 3**
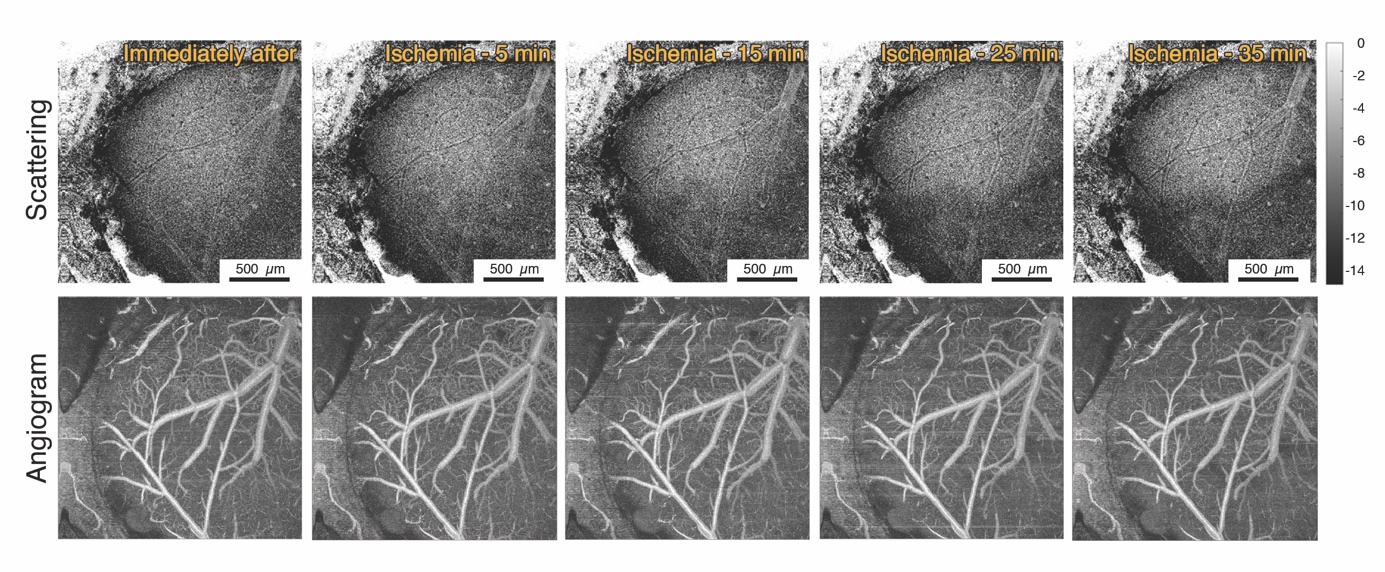

Abnormal focal attenuation changes focally appeared in the ischemic core area ~15-25 min after initiation of dMCAO while capillary perfusion immediately affected within the same zone. Scale bars: 500 µm. Unit of calibration bars is 1/mm^2^.

**Supplementary Figure 4**

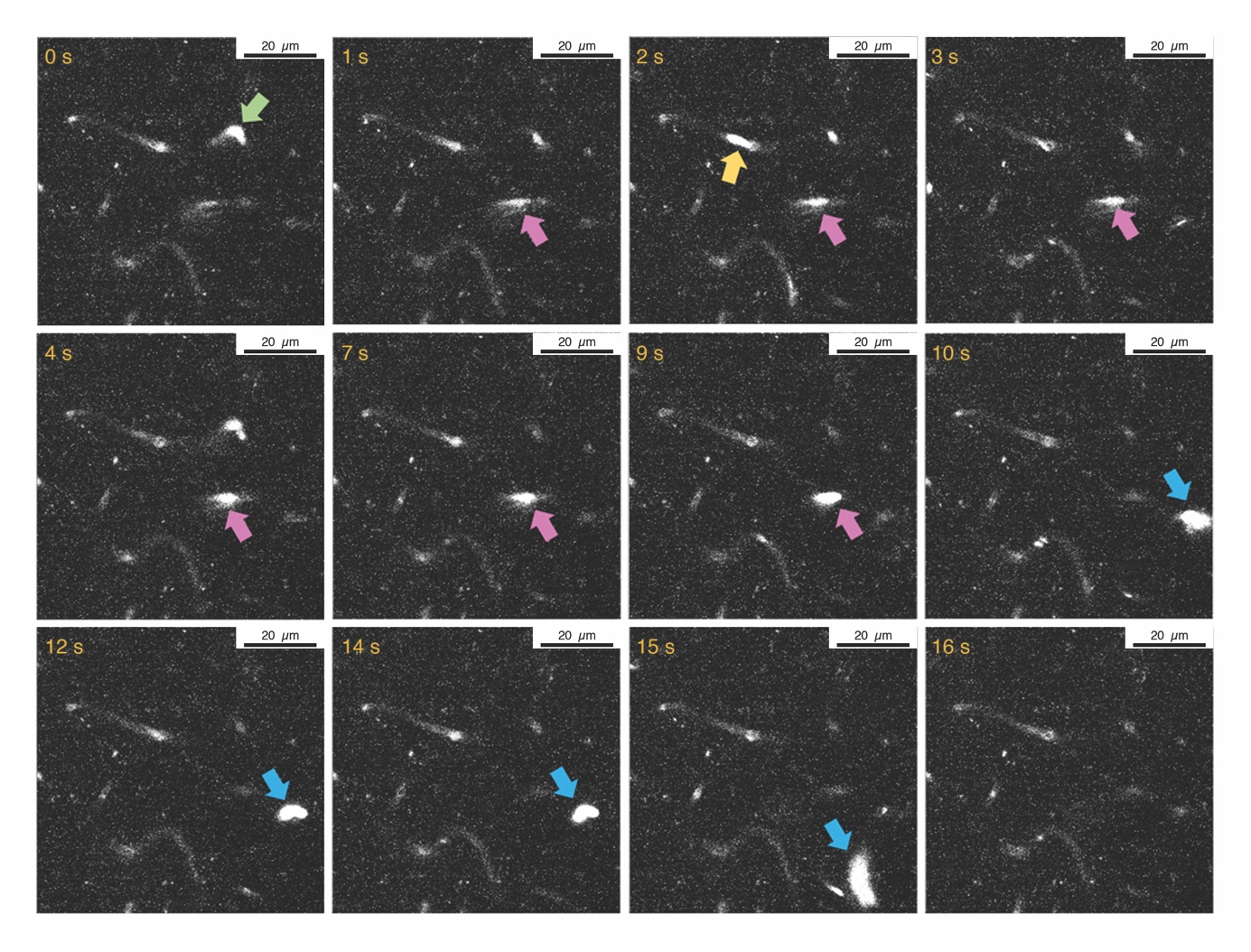

Two-photon microscopy time series of fluorescent leukocytes dynamically getting stuck in capillary network. Each particular leukocyte is indicated with different colors. Scale bar: 20 µm.

**SUPPLEMENTARY MOVIES**

**Supplementary Movie 1**

Laser speckle contrast imaging during baseline and dMCAO. An initial drop in MCA flow is subsequently followed by a periinfarct depolarization wave. Distal pial branches of MCA are supplied by retrograde flow from collateral vessels during occlusion. Darker colors indicate higher flow.

**Supplementary Movie 2**

Laser speckle contrast imaging during reperfusion after 60 min of dMCAO. Recanalization is confirmed by the rapid increase in MCA flow (darkening of color) accompanied by diminished collateral flow.

**Supplementary Movie 3**

Temporally averaged capillary angiograms do not provide the information on the flow dynamics of individual capillaries, i.e. whether they are flowing with RBCs continuously or flow is being interrupted (stalled) temporarily or permanently. OCT angiogram time series showing a 600x600 µm area for 6.5 minutes show that most capillaries approximately 2 hours after reperfusion in the ischemic penumbra experience dynamically interrupted flow; RBC motion spontaneously ceases and restarts. When a particular segment is has RBC flow, it is visible in white, and when RBC flow stops, the segment disappears in the image.

**Supplementary Movie 4**

OCT angiogram time series showing a 600x600 µm area in the ischemic penumbra during baseline, MCAO and recanalization. Stalling capillary segments can be easily recognized as they disappear and reappear with fluctuations in RBC flow. Interframe interval is 6.5 seconds. Note the very high frequency of stalls during ischemia but also persistently for 2 hours after recanalization, compared to baseline conditions.

**Supplementary Movie 5**

Two-photon fluorescent angiogram time series in ischemic penumbra 2 hours after reperfusion disclosed repetitive plugging of capillary flow with cells identified as leukocytes considering their large size (white arrows) compared to numerous red blood cells flowing. We were able to document both flow cessation and reinitiation events as well as moments of leukocytes squeezing through stalled capillaries.

**Supplementary Movie 6**

In these two-photon fluorescent angiogram time series in ischemic penumbra, repeated transient plugging of a capillary segment by squeezing leukocytes (arrow) affects the flow distribution in the immediately connected capillary branches, causing RBC flow in those segments to stall as well.

**Supplementary Movie 7**

Time series of fluorescent leukocytes labeled with Rhodamine 6G. As they enter and stay briefly (marked with arrows) in the imaging plane at 150 µm cortical depth, leukocytes can be visualized as globular circular or ovoid cells. In this video acquired from an isotype-control antibody injected animal, at 2 hours after reperfusion, we can see an accumulation of stagnant leukocytes in the microcirculatory bed.
